# A robust life-or-death selection platform for enzyme evolution

**DOI:** 10.1101/2023.10.09.561342

**Authors:** Suzanne C. Jansen, Clemens Mayer

**Affiliations:** Biomolecular Chemistry & Catalysis, Stratingh Institute, University of Groningen, Nijenborgh 4, 9747 AG Groningen, The Netherlands

## Abstract

Life-or-death selections evaluate the fitness of individual organisms on a population level. In enzyme engineering, such growth selections allow the rapid and straightforward identification of highly efficient biocatalysts from extensive libraries. However, selection-based improvement of (industrially-relevant) biocatalysts is challenging, as they require highly dependable strategies that artificially link their activities to host survival. Here, we showcase a robust and scalable life-or-death selection platform centered around the complementation of non-canonical amino acid-dependent bacteria. Specifically, we demonstrate how serial passaging of populations featuring millions of carbamoylase variants autonomously selects biocatalysts with up to 90,000-fold higher initial rates. Notably, selection of replicate populations enriched diverse biocatalysts, which feature distinct amino-acid motifs that drastically boost carbamoylase activity. As beneficial substitutions also originated from unintended copying errors during library preparation or cell division, we anticipate that our life-or-death selection platform will be applicable to the continuous, autonomous evolution of diverse biocatalysts in the future.

## Introduction

Evolution is a dynamic, all-purpose problem solver that enzyme engineers mimic in the laboratory to generate tailor-made biocatalysts.^1-4^In general, such directed evolution campaigns rely on four principal steps: (1) generating genetic diversity, (2) translating the genetic information into biocatalysts, (3) identifying variants with desired properties, and (4) recovering genes encoding improved enzymes (**Fig. 1A**).^5-7^While concurrent molecular biology tools readily enable the creation, translation, and recovery of vast gene libraries, exhaustive sampling of the resulting sequence space remains a persistent bottleneck for most directed evolution campaigns.^8,9^Therefore, general strategies to identify improved biocatalysts are highly sought-after. Ideally, such strategies should not only be fast, accurate, cheap, and easy to implement, but also allow for a high throughput and have the ability to differentiate enzymes with vastly different activities (=dynamic range, **Fig. 1B**).

**Figure 1:**
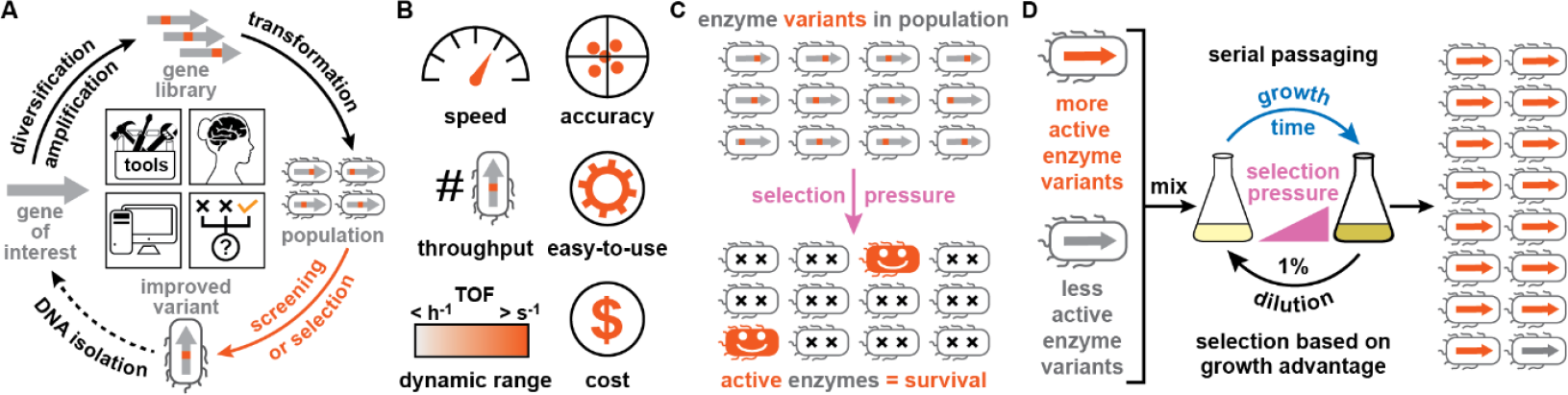
Directed evolution of biocatalysts by life-or-death selections. **A:** To tailor a gene of interest, directed evolution approaches rely on iterative cycles of genetic diversification, translation, and identification and isolation of improved enzyme variants. **B:** Strategies to identify improved biocatalysts should ideally meet a broad set of criteria. **C:** Selections assess all enzyme variants in a population simultaneously and eliminate undesired variants with an applied selection pressure. **D:** Serial passaging under selection pressure can amplify more active enzyme variants in a mixed population by virtue of the growth advantage they bestow on their host. Panel **C** was adapted from Ref. 16.

In principle, life-or-death selections have the potential to meet all these criteria.^10-12^First, they can rapidly assess all library members of vast populations simultaneously by applying a selective pressure that effectively eliminates undesired enzyme variants (**Fig. 1C**). Moreover, life-or-death selections are cheap and straightforward to implement in the laboratory, as identifying improved biocatalysts typically involves either plating populations on selective media or serially (or continuously) culturing populations under selective conditions. In the latter scenario, more active biocatalysts will gradually outcompete less proficient ones (**Fig. 1D**).^13-16^Lastly, previously employed selections for biocatalysts that directly confer a growth advantage to a host organism (e.g. chorismate mutase or β-lactamase) could be employed to identify enzyme variants with a wide range of activities.^17-19^

Despite these advantages, life-or-death selections are rarely employed for the directed evolution of (industrially-relevant) enzymes, as such biocatalysts typically do not confer a growth advantage to a producing organism. Instead, the use of (synthetic) auxotrophs,^20,21^genetic circuits,^22,23^or other sensors^24,25^is necessary to artificially link the activity of a desired biocatalyst to cell fitness/survival. While such chemical *complementation systems* are effective to interrogate individual enzyme variants of small, controlled populations, they often lack the robustness and/or scalability to elicit highly active biocatalysts from gene libraries with millions of members. For example, genetically-modified host cells are prone to escape their confinement by genomic mutations or other, unintended mechanisms, thereby increasing false-positive rates during selections.^26,27^Moreover, if products of an enzymatic transformation readily diffuse into the media, growth advantages are not confined to library members with high activities, but also enable non-beneficial ‘hitchhikers’ to remain in the population.^28^

To facilitate a more widespread use of life-or-death selections for the directed evolution of biocatalysts, we have recently showcased a chemical complementation strategy that leverages recoded organisms addicted to non-canonical amino acids (ncAAs, *vide infra*).^16^Here, we demonstrate the scalability of our approach and its ability to autonomously elicit a diverse panel of efficient biocatalysts from highly diverse populations. Excitingly, serial passaging proved sufficient to amplify inherently ‘scarce’ variants within these populations, including enzyme variants whose improved performance results from unintended, random mutations that were introduced during PCR cycles or cell division. Combined, these results showcase not merely the impressive potential and simplicity of our life-or-death selection for enzyme engineering, but also encourage its application in autonomous exploration of extended evolutionary trajectories of mechanistically-diverse biocatalysts in the future.

## Results and Discussion

### Selecting improved enzymes from heavily-skewed populations

For our recently reported chemical complementation strategy, we first created bacteria, whose lives in presence of carbenicillin were dependent on the supply and incorporation of a non-canonical amino acid (ncAA) into an engineered β-lactamase.^29,30^Next, we leveraged the ncAA-dependency of such *recoded bacteria* to evolve a biocatalyst that could yield these ncAAs from externally-provided, synthetic precursors (**Fig. 2A**).^16^As proof-of-principle, we applied our platform to engineer a *L*-N-carbamoylase from *Sinorhizobium meliloti* strain CECT 4114, SmLcar,^31^to promote the hydrolysis of the *N*-carbamoylated ncAA, cam-*L*-3-nitro-tyrosine (cam-3nY, **Fig. 2B**). We found that growth rates of *E. coli* cells in presence of carbenicillin correlated with the activities of produced SmLcar variants. Sequentially randomizing positions Leu217 and Phe329 enabled us to identify SmLcar_GY (named after its Leu217Gly and Phe329Tyr substitutions), which displayed shorter lag-times under selective conditions and proved >1,000-times more proficient than the wildtype carbamoylase.^16^Lastly, we also demonstrated that serial passaging could faithfully elicit SmLcar_GY from a small, controlled population containing 20 enzyme variants.^16^

**Figure 2:**
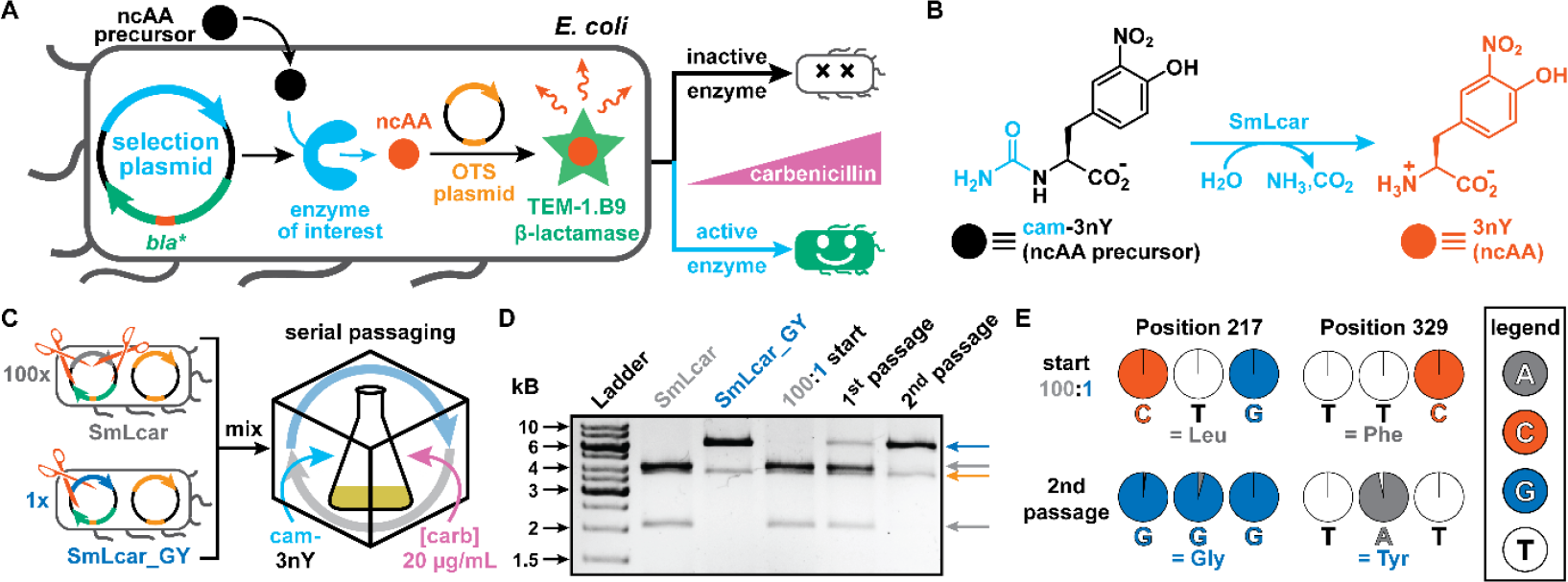
Selecting improved variants from heavily-skewed populations. **A:** Blueprint of a life-or-death selection platform based on chemical complementation. An enzyme of interest converts a synthetic precursor to a non-canonical amino acid (ncAA), which is incorporated into an engineered β-lactamase using an orthogonal translation system (OTS) that suppresses an in-frame stop-codon. Active enzymes enable growth of *E. coli* in presence of carbenicillin, while bacteria featuring inactive ones perish under the selection pressure. **B:** The carbamoylase SmLcar catalyzes the hydrolysis of cam-3nY and gives rise to the ncAA *L*-3-nitro-tyrosine (3nY). **C:** Modified selection plasmids with restriction sites allow visual differentiation of SmLcar and SmLcar_GY in a skewed population during serial passaging. **D:** Restriction digest followed by agarose gel electrophoresis reveals quantitative patterns for SmLcar and SmLcar_GY during serial passaging. The band at 6085 bp (blue arrow) depicts a linearized plasmid for SmLcar_GY (cut once). The bands at 4016 bp and 2069 bp (grey arrows) depict two fragments of the SmLcar plasmid (cut twice). The band at 3698 bp (orange arrow) represents the OTS plasmid, which is present in both. **E:** The relative base distribution for codons at position 217 and 329 during the mock selection, obtained from Sanger sequencing pooled plasmids of the mixed populations. Panels **A-B** were adapted from Ref. 16.

With the aim of selecting efficient biocatalysts from large libraries, we first set out to gain quantitative insights into the amplification of improved enzyme variants in a heavily-skewed population. Toward this end, we constructed selection plasmids, which allow for a straightforward differentiation of SmLcar and SmLcar_GY following restriction digest. In brief, strategically placed restriction sites yield two fragments for SmLcar upon digestion, while only linearizing SmLcar_GY (**Fig. 2C**, see **Supplementary Information** for details). With these constructs, we performed a mock selection by mixing cultures containing SmLcar and SmLcar_GY in a 100:1 ratio. This heavily-skewed population was subjected to serial passaging under selective conditions – that is cultures were grown for ∼24 hours in presence of 20 μg/mL carbenicillin and 500 μM cam-3nY, followed by 100-fold dilutions in fresh selective medium. Performing restriction digests on isolated plasmids from the starting and selected populations revealed that selection consistently favors amplification of SmLcar_GY (**Fig. 2D**). In fact, two serial passages sufficed for SmLcar_GY to become dominant. Subjecting the isolated plasmids to Sanger sequencing independently confirmed that the population was essentially devoid of the wildtype carbamoylase following two serial passages (**Fig. 2E**). Combined, these results indicate that the growth advantage more active biocatalysts provide suffices to reliably amplify them, even when starting at a considerable disadvantage.

### Design, preparation, and evaluation of SmLcar libraries

After successful enrichment of SmLcar_GY in the mock selection, we set out to retrieve highly active SmLcar variants from increasingly diverse populations. Toward this end, we designed two SmLcar libraries, Lib_2N and Lib_4N, by either randomizing two or four residues that line the binding pocket of SmLcar (**Fig. 3A**). Specifically, Lib_2N simultaneously targets Leu217 and Phe329, which have shown high synergy when sequentially randomized in our previous work that elicited SmLcar_GY.^16^Evaluating all 400 possible amino-acid combinations at once should thus shed further light on potential epistatic interactions between these two residues. Lib_4N additionally includes the neighboring residues Gln215 and Gly327, whose amino acid side chains also project toward the binding pocket (**Fig. 3A**). Drastically increasing the genetic diversity from ≈10^3^for Lib_2N to ≈10^6^in Lib_4N should enable us to uncover multiple, new “solutions” toward boosting the activity of SmLcar for the hydrolysis of cam-3nY. Following standard procedures for the preparation of Lib_2N and Lib_4N, we obtained 2.4×10^3^transformants for Lib_2N and 8.4×10^6^transformants for Lib_4N, thus exceeding the theoretical genetic diversity of these libraries by a factor of 2.4 and 8.4, respectively (**Supplementary Table S1**).

**Figure 3:**
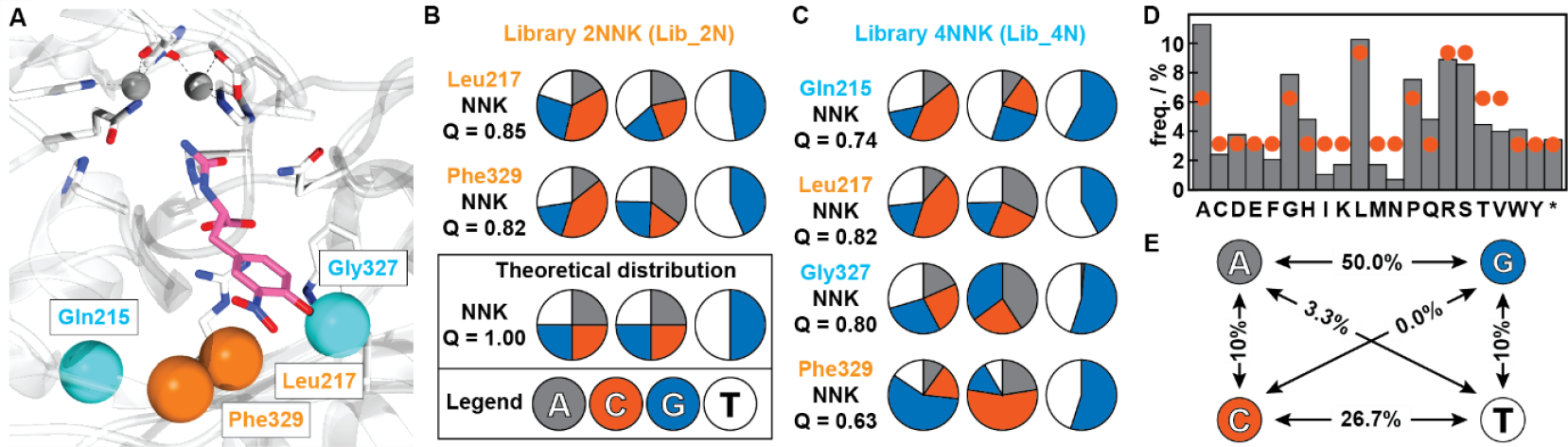
Library design and quality. **A:** Close-up view of the active site of SmLcar (PDB: 8APZ). Orange and cyan spheres indicate positions targeted for site-directed mutagenesis. Two bivalent metal ions and residues critical for their binding are given, and the substrate cam-3nY (pink) is docked in the active site. **B-C:** Pie charts based on base calls from Sanger sequencing of Lib_2N populations (**B**) or 71 individual Lib_4N library members (**C**) depict the relative distribution of bases at each targeted position. Q-values are given as a measure of diversity. The theoretical distribution is based on the perfect NNK codon and has a Q-value of 1.00. **D:** Bar chart depicting the observed distribution of amino acids for Lib_4N (grey bars) and expected theoretical distribution (red dots). **E:** The percentage of transitions (left/right) and transversions (up/down, diagonal) observed in Lib_4N based on 30 random mutations found in sequence data of 71 library members in Lib_4N.

Next, we introduced the libraries into our recoded bacteria and examined their composition by subjecting either pooled populations (Lib_2N) or 71 individual library members (Lib_4N) to Sanger sequencing (see **Supplementary Information** for details). Calculating the Q-values^32^of each NNK stretch, which is a quantitative analysis of library degeneracy, revealed a high degree of genetic diversity for both libraries (**Figs. 3B-C** and **Supplementary Table S2**). Specifically, Lib_2N displays Q-values of 0.85 and 0.82, indicating a near-equal distribution of the desired nucleotides at both NNK stretches. While by comparison Lib_4N scores slightly worse (Q-values ranging from 0.63-0.82), the genetic diversity across all four targeted positions gives rise to a distribution of amino acids that closely tracks the frequencies expected for NNK codons (**Fig. 3D**). Notably, we were unable to detect thymidine at the second randomized nucleotide in position Gly327 in any of the inspected library members (**Fig. 3C**), reducing the diversity of Lib_4N by 19%.

At the same time, we surmised that relying on three PCRs to amplify (part of) the SmLcar gene for library preparation could have inadvertently increased the diversity of Lib_2N and Lib_4N by accumulating random copying errors. Indeed, when analyzing high quality sequence data of 71 individual Lib_4N variants (**Supplementary Data File 1**), we identified on average 0.71 random mutations per SmLcar gene. This number is consistent with the low error rate of the employed high-fidelity DNA polymerase (≈1×10^-6^substitutions per base). The majority (77%) of these mutations were transition substitutions (50% for A ⇄G and 27% for C ⇄T), which involve interchanges of purines or pyrimidines (**Fig. 3E** and **Supplementary Table S3**). Critically, these random mutations add to the overall genetic diversity of the libraries, with variants featuring beneficial amino acid substitutions having the chance to be amplified over consecutive serial passages. Taken together, these results thus attest that Lib_2N and Lib_4N display sufficient genetic diversity, which results from randomizing key residues lining the binding pocket of SmLcar and random mutations introduced during library preparation.

### Autonomous selection of proficient carbamoylases from large libraries

With sufficiently diverse libraries in hand, we next challenged our life-or-death selection to autonomously elicit a panel of highly active SmLcar variants. Following library generation and transformation into recoded hosts, the selection of proficient carbamoylase based on growth advantage is as follows: (1) serial passaging of replicate populations, while increasing the selection pressure from mild to moderate (20, to 50, to 100 μg/mL carbenicillin); (2) plating of enriched variants on LB agar; (3) evaluation of single colonies’ growth rate in the plate reader under moderate or high selection pressure (100 μg/mL or 500 μg/mL carbenicillin), and (4) identifying SmLcar variants that grew fasted at the high selection pressure by sequencing (see **Supplementary Information** for details). To probe whether the selection of proficient SmLcar variants is subject to stochasticity, we performed serial passaging for two Lib_2N populations (2N_A-B) and four Lib_4N populations (4N_A-D) in parallel (**Fig. 4A**). With the exception of 4N_B, which went extinct following the second serial passage, the performed selections thus bestowed on each surviving population a unique *evolutionary history* (**Extended Data Fig. 1**).

**Figure 4:**
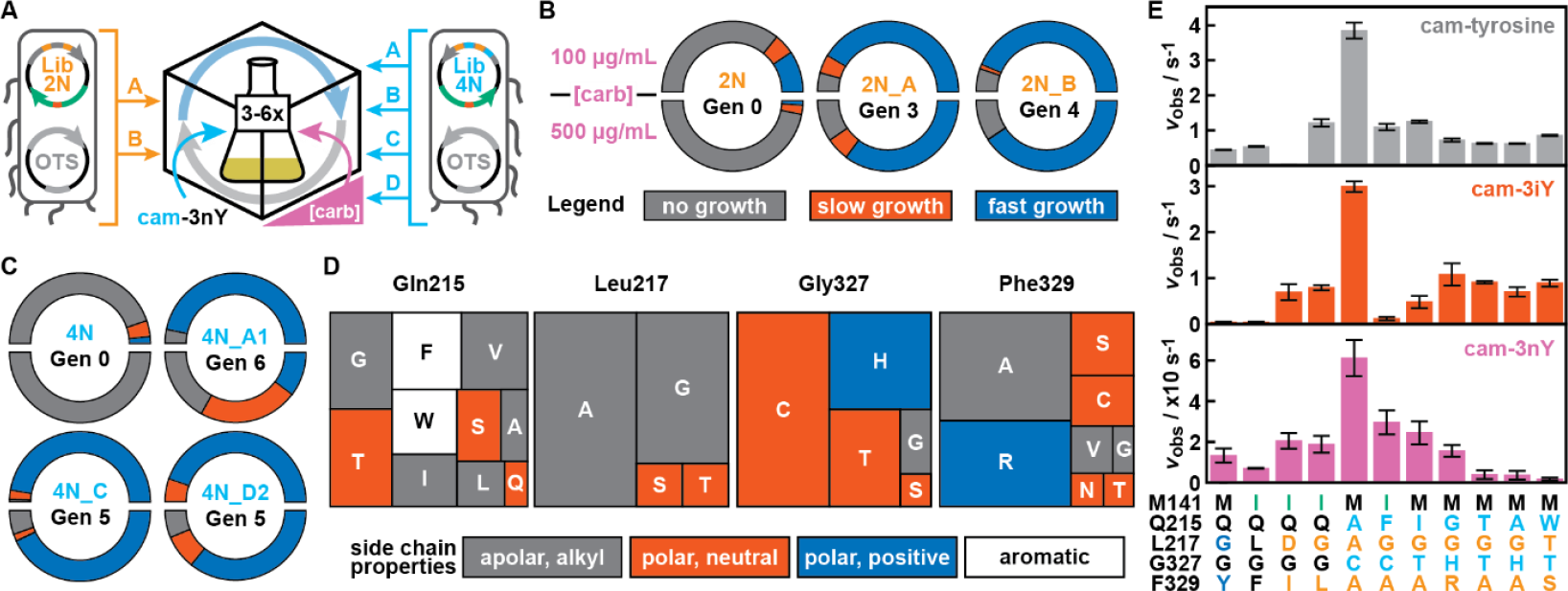
Autonomous selection of proficient SmLcar variants. **A:** Workflow for the identification of improved SmLcar variants from highly diverse populations by serial passaging. **B-C:** Radial bar charts depicting the distribution in growth phenotypes for start and endpoints of 2N populations (**B**) and 4N populations (**C**). Distributions follow from growth under 100 μg/mL (upper half) or 500 μg/mL (lower half) carbenicillin pressure. No growth is defined as failing to exceed an OD_600_ of 0.3 within 48 hours; slow growth is defined as exceeding an OD_600_ of 0.3 after 24 to 48 hours; fast growth is defined as exceeding an OD_600_ of 0.3 within 24 hours. **D:** Tree map charts depicting the diversity within amino acids across all four randomized positions in 132 evaluated SmLcar variants (37 unique variants) that bestowed a fast-growing phenotype. **E:** The initial rate (v_obs_ in s^-1^) of reference variant SmLcar_GY and the panel of evolved SmLcar variants for (nc)AA precursors cam-3nY, cam-3iY, and cam-Y. Reactions were performed in triplicates at 25 °C. The corresponding parameters are listed in Table 1.

For populations 2N_A and 2N_B, we performed three and four serial passages, respectively, at which point selections had curtailed the genetic diversity of these populations to a few genotypes as judged by Sanger sequencing (**Extended Data Fig. 2**). As anticipated, the global fitness of the population steadily increased with each passage, as judged by faster growth (=shorter lag times) of aliquots taken after each growth cycle (**Extended Data Fig. 2**). To quantify this fitness increase, we compared the growth characteristics of 95 individual colonies taken from the start and 95 individual colonies taken from the respective endpoints of these populations (**Fig. 4B** and **Supplementary Table S4**). When subjected to the moderate selection pressure (=100 μg/mL carbenicillin) present at the end of the serial passaging regime, 19% of the colonies taken from the initial populations displayed a fast-growing phenotype – defined as exceeding an OD_600_ of 0.3 within 24 hours or less (**Fig. 4B**). This fraction was markedly increased in the selected populations, with 83% and 88% of all single colonies being categorized as fast-growing in 2N_A and 2N_B, respectively. To identify bacteria featuring the most active SmLcar variants, we challenged all variants to grow at a high selection pressure (=500 μg/mL carbenicillin). The difference between starting and selected populations became more pronounced, as only 2% of the initial library members but 70% and 81% of 2N_A and 2N_B displayed fast growth (**Fig. 4B**).

**Table 1:** The kinetic parameters of SmLcar and evolved variants for (nc)AA precursors cam-3nY, cam-3iY, and cam-Y were determined at pH 6 for cam-3nY and pH 8 for cam-3iY and cam-Y. Reactions were performed in triplicate at 25 °C, with 2 mM substrate and 1 mM MnCl2. Average v_obs_ is given, with standard deviation in brackets. An asterisk (*) indicates SmLcar variants from 2N and 4N libraries harboring the Met141Ile substitution.

| Enzyme | cam-L-3-nitrotyrosine<br>(cam-3nY) |  | cam-L-3-iodotyrosine<br>(cam-3iY) |  | cam-L-tyrosine<br>(cam-Y) |  |
| --- | --- | --- | --- | --- | --- | --- |
| | $v_{\text{obs}}$ [s<sup>-1</sup>] | relative | $v_{\text{obs}}$ [s<sup>-1</sup>] | relative | $v_{\text{obs}}$ [s<sup>-1</sup>] | relative |
| SmLcar | $7.87 \times 10^{-5}$<br>($1.66 \times 10^{-5}$) | 1 | $3.33 \times 10^{-5}$<br>($1.10 \times 10^{-6}$) | 1 | 0.012<br>($2.59 \times 10^{-4}$) | 1 |
| SmLcar_GY | 0.13<br>(0.034) | 1,690 | 0.040<br>(0.009) | 1,210 | 0.44<br>(0.009) | 37 |
| SmLcar_M141I | 0.069<br>(0.002) | 880 | 0.039<br>(0.0010) | 1,160 | 0.52<br>(0.0088) | 44 |
| SmLcar_DI* | 0.20<br>(0.038) | 2,600 | 0.69<br>(0.18) | 20,800 | 0.023<br>(0.0036) | 2 |
| SmLcar_GL* | 0.19<br>(0.042) | 2,390 | 0.79<br>(0.055) | 23,700 | 1.21<br>(0.11) | 103 |
| SmLcar_AACA | 0.61<br>(0.090) | 7,760 | 2.99<br>(0.12) | 89,800 | 3.85<br>(0.23) | 326 |
| SmLcar_FGCA* | 0.30<br>(0.059) | 3,760 | 0.12<br>(0.037) | 3,500 | 1.09<br>(0.094) | 93 |
| SmLcar_IGTA | 0.24<br>(0.056) | 3,100 | 0.48<br>(0.13) | 14,300 | 1.25<br>(0.039) | 106 |
| SmLcar_GGHR | 0.16<br>(0.030) | 1,980 | 1.08<br>(0.24) | 32,300 | 0.72<br>(0.048) | 61 |
| SmLcar_TGTA | 0.039<br>(0.021) | 500 | 0.91<br>(0.026) | 27,300 | 0.63<br>(0.017) | 53 |
| SmLcar_AGHA | 0.036<br>(0.021) | 460 | 0.70<br>(0.10) | 21,000 | 0.62<br>(0.012) | 53 |
| SmLcar_WTTS | 0.017<br>(0.007) | 220 | 0.89<br>(0.077) | 26,700 | 0.86<br>(0.015) | 73 |

Following manual inspection of individual growth curves, we selected the 10 best-performing colonies of 2N_A and 2N_B and identified the SmLcar variants they harbored by sequencing. Consistent with our previous evolutionary campaign, 2N_A had converged to SmLcar_GY variants, which all featured the previously identified Leu217Gly and Phe329Tyr substitutions, but differed by (silent) random mutations in their genes (**Supplementary Table S5**). Strikingly, sequencing the best-performing carbamoylase variants from 2N_B revealed markedly different genotypes. While SmLcar_GY was only identified once, the other carbamoylase variants featured six times an Asp217/Ile329 combination (SmLcar_DI) and three times Gly217/Leu329 (SmLcar_GL). Upon closer inspection, both SmLcar_DI and SmLcar_GL shared an additional random Met141Ile substitution (**Supplementary Table S5)**, which was outside of the targeted positions but affected a residue that that also lines the binding pocket (**Extended Data Fig. 3**). Since the otherwise dissimilar variants shared this substitution, we hypothesized that Met141Ile by itself could be a beneficial mutation that provides a growth advantage. To investigate this hypothesis, we prepared SmLcar_M141I and observed that this variant indeed provided a fast-growing phenotype to producing bacteria at 100 μg/mL carbenicillin, but a slow-growing phenotype at 500 μg/mL carbenicillin (**Extended Data Fig. 3**). To confirm whether shorter lag times observed from selected variants translate to higher catalytic activities, we chose SmLcar_DI, SmLcar_GL, and SmLcar_M141I for further kinetic characterization (*vide infra*).

For populations 4N_A, C, and D, we performed 5 to 6 serial passages before examining their composition and the growth characteristics of individual colonies. Sanger sequencing of pooled plasmids revealed that selections had curtailed the diversity of initial libraries (**Extended Data Fig. 4**), with 4N_A and 4N_C showing the apparent enrichment of a handful of phenotypes. Evaluating the growth curves for ∼30 colonies of 4N_A-D attested on the successful selection of fast-growing phenotypes at a moderate selection pressure (100 μg/ml), but also highlighted that 4N_A and 4N_D contained only a small number of variants that could grow rapidly at 500 μg/ml (**Supplementary Table S4**). In an attempt to exploit the stochastic nature of selections to improve the fitness of 4N_A and 4N_D, we revived glycerol stocks from their second passages and performed an additional three selective growth-dilution cycles to yield 4N_A2 and 4N_D2 (**Extended Data Fig. 1**). While these measures did not prove successful for the former, population 4N_D2 had enriched variants that displayed fast growth at the high selection pressure. (**Supplementary Table S4**).

To compare the apparent fitness increase of all 4N populations, we determined the growth characteristics of 95 individual colonies taken from the start and 349 individual colonies taken from the respective endpoints, i.e. 4N_A1, 4N_C, and 4N_D2 (**Fig. 4C** and **Supplementary Table S4**). Only 3% of colonies taken from the initial populations displayed a fast-growing phenotype when subjected to the moderate selection pressure (100 μg/mL carbenicillin) that is present at the end of the serial passaging regime (**Fig. 4C**). Conversely, our selections successfully enriched fast-growing phenotypes, with 94%, 95%, and 89% of colonies in 4N_A1, 4N_C, and 4N_D2, respectively, exceeding an OD_600_ of 0.3 in under 24 hours. Substantially increasing the stringency to 500 μg/mL carbenicillin halted proliferation entirely in the initial population, while 20% of 4N_A1, 83% of 4N_C, and 72% of 4N_D2 colonies still displayed a fast-growing phenotype (**Fig. 4C**). Taken together, these results showcase that serial passages do not only effectively discriminate host fitness levels at the applied pressure, but also suffice to amplify variants that can thrive under higher selection pressure.

To pinpoint genotypes that were enriched during selections of 4N libraries, we compared sequencing data of the SmLcar genes for 73 single colonies prior to serial passages and of 132 fast-growing single colonies at the end of each evolution campaign. As expected, the initial population displayed diverse sequences at the randomized positions, with a good representation of each side chain property (**Extended Data Fig. 5** and **Supplementary Table S6**). Conversely, the diversity at these residues had greatly diminished for the 132 evaluated SmLcar variants that bestowed a fast-growing phenotype following growth-dilution cycles (**Fig. 4D** and **Supplementary Table S7**). Combined, the 37 unique sequences obtained offer key insights and commonalities on the type of SmLcar variants that were enriched across these populations (**Fig. 4D**). (1) Gln215 remained the most diverse position by accepting a variety of small, hydrophobic, aromatic, and polar residues, while being devoid of any charged residues. (2) For Leu217, variants featuring small side chains (Ala, Gly) became strongly enriched (89%). This finding is consistent with our previous evolutionary campaign, where we demonstrated that enlarging the binding pocket via a L217G substitution is critical to accommodate cam-3nY in a productive conformation, in which the aromatic side chain is flipped by ∼90°.^16^(3) The substitutions at Gly327 and Phe329 reveal two distinct *solutions*, which are able to drastically boost the growth rates of recoded bacteria (**Fig. 4D**). In one solution, a small hydrophobic or polar residue at position Phe329 is paired with a residue at Gly327 that provides a hydrogen-bond donor at Cβ (i.e. Cys, Ser, or Thr). In the alternative solution, SmLcar variants feature a His at position 327, which is almost exclusively combined with a positively-charged Arg at position 329.

### In vitro characterization of improved SmLcar variants

Based on the observed genotypes in selected 2N and 4N libraries, we chose a total of 10 SmLcar variants that granted a fast-growing phenotype for further characterization (**Table 1, Supplementary Tables S5** and **S7**). In brief, these are SmLcar_M141I as well as SmLcar_DI and SmLcar_GL, which were identified in 2N_B and also harbored this apparently beneficial yet unintended Met141Ile substitution. From the 4N populations, we included the most amplified variant from each surviving population, which are SmLcar_WTTS (4N_A1), SmLcar_IGTA (4N_C), and SmLcar_TGTA (4N_D2). Additionally, we selected representatives for the two solutions identified in the sequencing analysis, namely SmLcar_AACA and SmLcar_GGHR, as well as a hybrid variant, SmLcar_AGHA, which was the only identified variant that features His327 but *not* Arg329. Lastly, we chose SmLcar_FGCA, whose substitutions not only follow the first motif, but which also features the randomly introduced Met141Ile substitution observed in 2N_B variants. Lastly, we included the wildtype and the previously described SmLcar_GY as points of reference for a low and high enzyme activity.

Prior to kinetic characterization, we aimed to eliminate the possibility that the fitness increase in hosts harboring evolved SmLcar variants had resulted from undesirable adaptations or escapes. Toward this end, we cloned the SmLcar genes of our panel anew and transformed them into fresh recoded hosts. Manual inspection of individual growth curves showed that even at 500 μg/mL carbenicillin, all evolved variants still presented short lag times of ∼6–12 hours, in accordance with the reference variant SmLcar_GY (**Extended Data Fig. 6**). Critically, the absence of escapes or false positives affirms the robustness of the genotype-phenotype linkage that is central to our life-or-death selection platform.

Next, to verify that we have elicited highly active biocatalysts, we purified SmLcar, SmLcar_GY, and all ten variants of the panel (**Supplementary Figure S1, Supplementary Table S8**), and determined their initial rates to hydrolyze three (nc)AA precursors (2 mM) in vitro (see **Table 1** and **Supplementary Information** for details). These are (1) cam-3nY, which was employed throughout the selection campaigns, (2) cam-3-*L*-iodo-tyrosine (cam-3iY), an alternative *m*-substituted tyrosine analog, and (3) cam-*L*-tyrosine (cam-Y). Consistent with previous reports,^16,31^SmLcar displays poor activity for all three substrates, but has a strong preference for cam-Y (*v*_obs_ = 0.71 min^-1^) over cam-3nY (0.28 h^-1^) and cam-3iY (0.12 h^-1^). Excitingly, all 10 SmLcar variants from our panel outperform the wildtype enzyme for all substrates – typically by 2-4 orders of magnitude depending on the carbamoylated precursor. For example, the most active variant SmLcar_AACA displays initial rates of 0.61 s^-1^for cam-3nY, 2.99 s^-1^for cam-3iY, and 3.85 s^-1^for cam-Y, thus exceeding wildtype activities by a factor of 7,760, 89,800, and 326, respectively (**Table 1** and **Fig. 4E**).

A comparison of the results obtained across all 10 SmLcar variants and three substrates provides some valuable insights (**Table 1** and **Fig. 4E**). (1) The selected carbamoylases are highly efficient catalysts, which outperform the previously evolved SmLcar_GY for at least two of the tested precursors. (2) We do not observe the selection of *specialists* that only display high activities for cam-3nY, the precursor used during serial passaging. In fact, SmLcar variants featuring substitutions TGTA, AGHA, and WTTS show significantly higher activities for cam-3iY and cam-Y than cam-3nY. (3) Conversely, we observe the selection of a number of *generalists*, which outperform SmLcar_GY for all three substrates. Specifically, SmLcar_GL, AACA, IGTA, and GGHR display comparable efficiencies on all substrates, suggesting that these substitutions present general catalytic solutions for the conversion of carbamoylated tyrosine and *m*-substituted analogs. (4) Lastly, the randomly introduced Met141Ile substitution is highly beneficial, with additional substitutions at the targeted positions in the 2N and 4N libraries further boosting its activity for cam-3nY. Variants featuring the Met141Ile substitution also show diverse reactivity profiles: SmLcar_FGCA maintains the reactivity profile of the random Met141Ile substitution, SmLcar_DI selectively converts *m*-substituted tyrosine analogs, and SmLcar_GL represents a generalist enzyme.

## Conclusion

Strategies to identify improved enzyme variants from large libraries are highly sought-after in the directed evolution of (industrially-useful) biocatalysts. Here, we demonstrated the robustness and scalability of a previously-described life-or-death selection platform by autonomously eliciting a diverse panel of efficient biocatalysts from large and diverse populations. Specifically, we used the chemical complementation of recoded hosts to investigate epistatic interactions of two or four simultaneously targeted positions within a carbamoylase. Despite starting from replicate populations, we observed the enrichment of carbamoylase variants with distinct substitutions – some of which were the result of random copying errors. Strikingly, each characterized carbamoylase outperformed the wildtype enzyme by at least two orders of magnitude, thus presenting a different “solution” to the same enzymatic “problem”.

In its current setup, this life-or-death selection platform already shows impressive potential for widespread application in enzyme engineering campaigns. First, our platform is fast, easy-to-implement, and requires no specialized and/or expensive equipment. Selections encompass serial passaging for ∼1 week under increasingly stringent conditions, with carbenicillin serving as tunable selection pressure and cell survival as common readout. Second, the platform is robust and high-throughput. We evolved highly efficient biocatalysts without encountering hitchhikers or cells that had escaped their confinement. With the largest library in this work containing >10^6^enzyme variants, its scalability appears only limited by transformation efficiency. Third, rather than enzymes specialized for the substrate used in selections, the platform elicits overall catalytic improvements. In fact, we retrieved a number of generalist enzymes that showed drastically increased rates on never-before-seen substrates. Fourth, our life-or-death selection strategy is flexible in nature. Centered around the enzyme-catalyzed production of ncAAs from synthetic precursors, the platform should be applicable to engineering mechanistically-diverse enzyme classes that can yield one of the >150 ncAAs that have been incorporated into proteins in vivo.^33^

Lastly, we anticipate that interfacing selections based on complementing recoded bacteria with gene-directed hypermutation strategies will result in the construction of a versatile continuous evolution platform.^34,35^In the future, such a system should enable us to navigate the sequence space of diverse biocatalysts autonomously and uncover highly proficient enzymes featuring exceedingly unpredictable mutations, which are far beyond our current rationalization.

## Acknowledgements

C.M. and S.C.J. are thankful to R. Rubini for the guidance with the chemical complementation workflow and to T.R. Oppewal for the help with library preparation. C.M. acknowledges the NWO (ENW-M grant OCENW.M20.278 and ENW-XS grant 01259958) for funding.

## 1. Extended Data Figures

**Extended Data Fig. 1:**
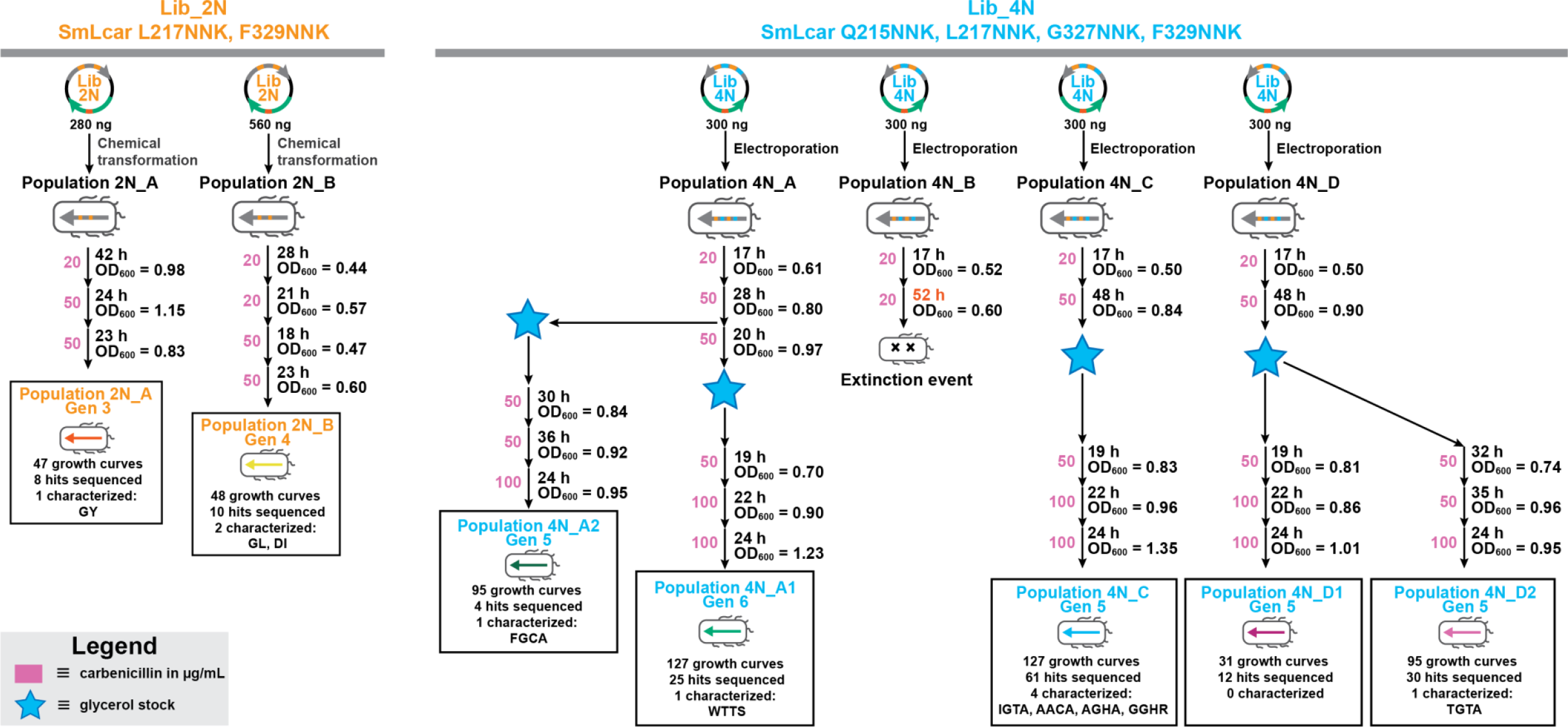
Evolutionary histories. A flowchart that represents each population’s unique evolutionary history. Arrows represent serial passages, and for each of them the scheme indicates selection stringency, incubation time and the end OD_600_. Arrows without these details represent non-selective precultures, such as those after reviving a glycerol stock.

**Extended Data Fig. 2:**
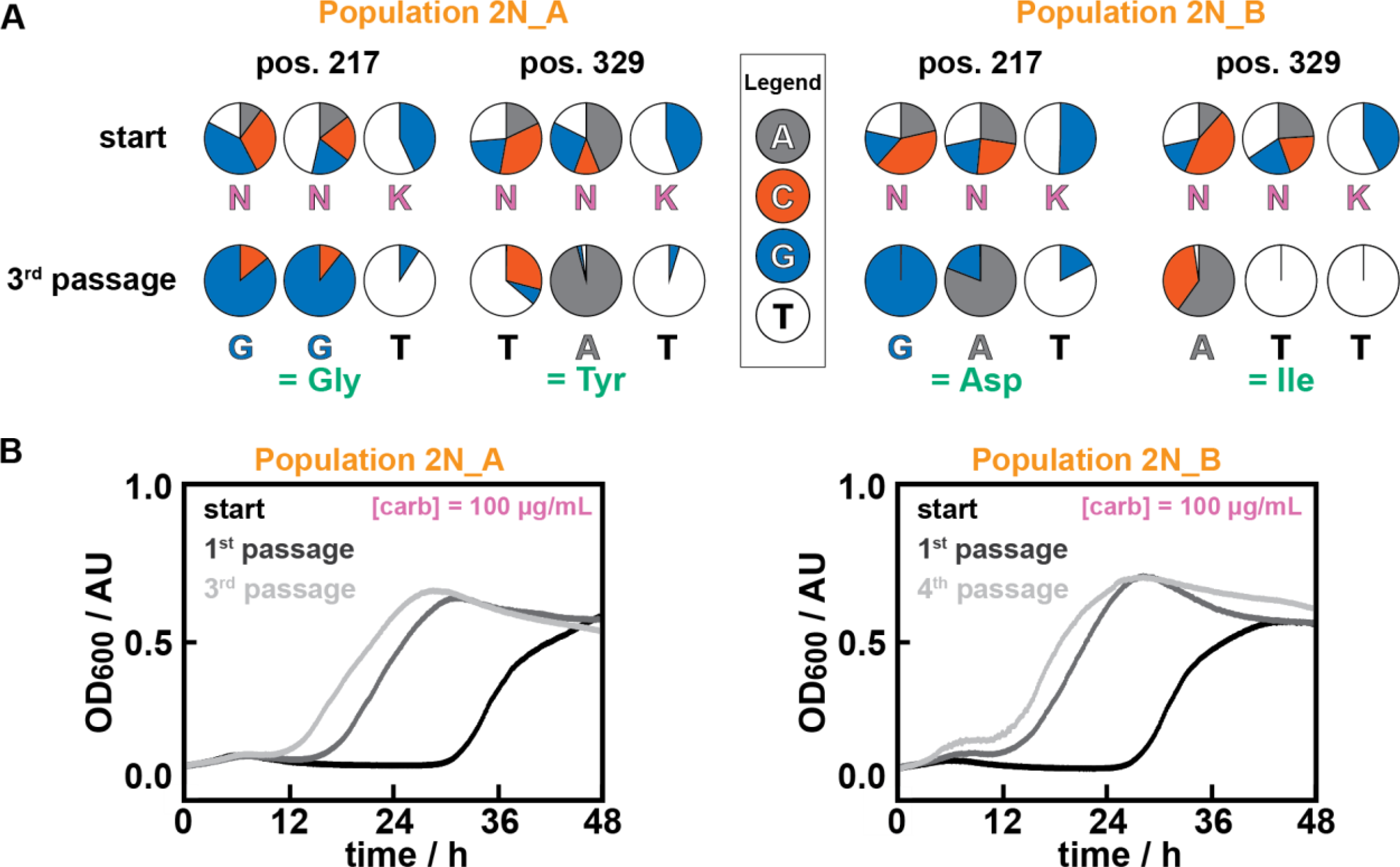
Genotypical and phenotypical changes in populations 2N_A and 2N_B during selection. **A:** The relative base distribution for codons at position 217 and 329 derived from base calls obtained from Sanger sequencing pooled plasmids. The consensus of the genotypes is given in green. **B:** Growth curves of the starting population, first, and final passage for population 2N_A and 2N_B. Note that shorter lag time indicate higher carbamoylase activity. Cells were grown in the presence of 500 μM cam-3nY.

**Extended Data Fig. 3:**
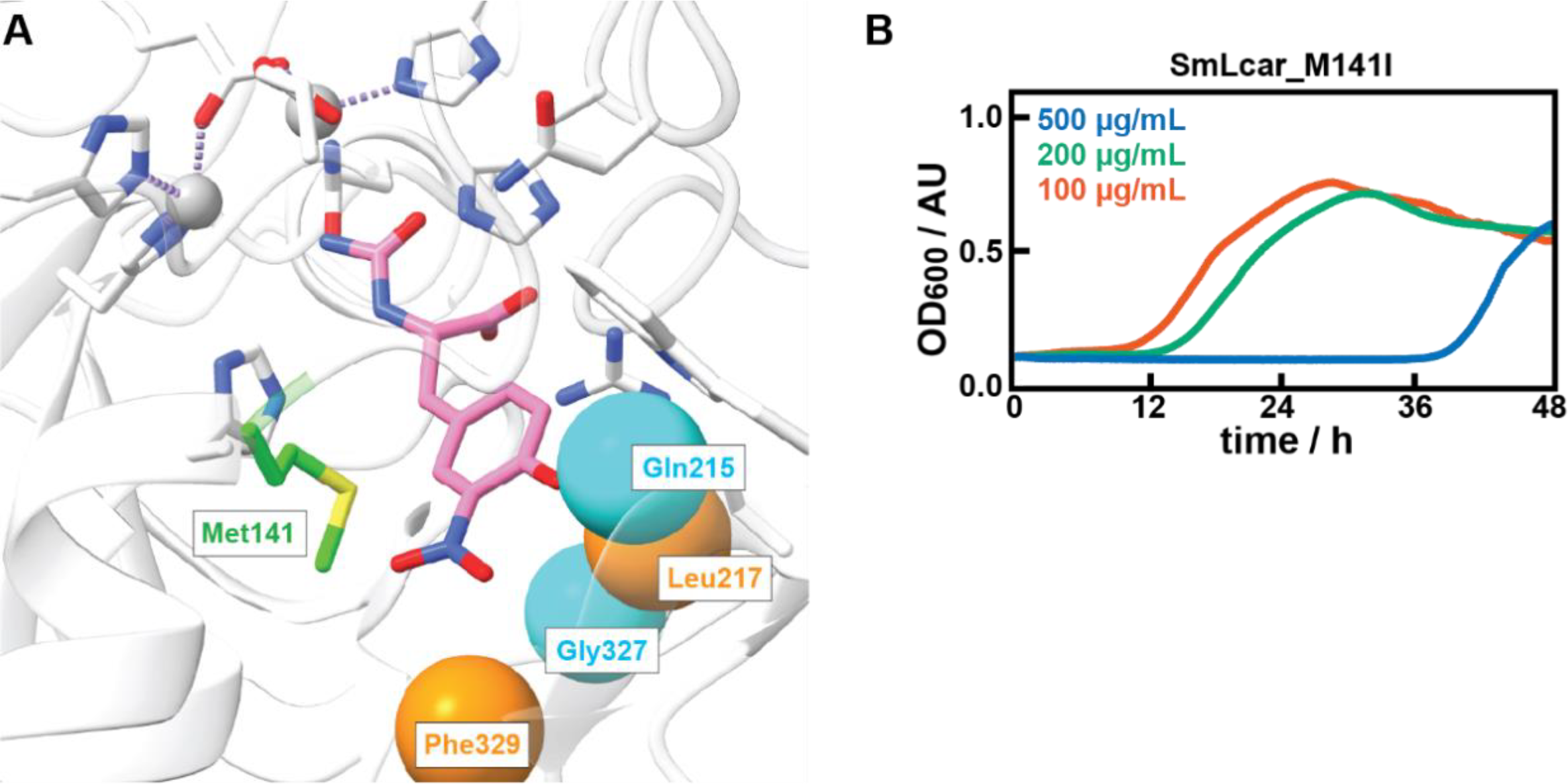
Analysis of random substitution Met141Ile. **A:** Close-up view of the active site of SmLcar (PDB: 8APZ) highlighting Met141 (green). The positions targeted by site-directed mutagenesis are also indicated (cyan and orange). The substrate cam-3nY (pink) is docked in the active site. **B:** Growth curves of the SmLcar_M141I variant in the presence of 500 μM cam-3nY at increasing carbenicillin concentrations.

**Extended Data Fig. 4:**
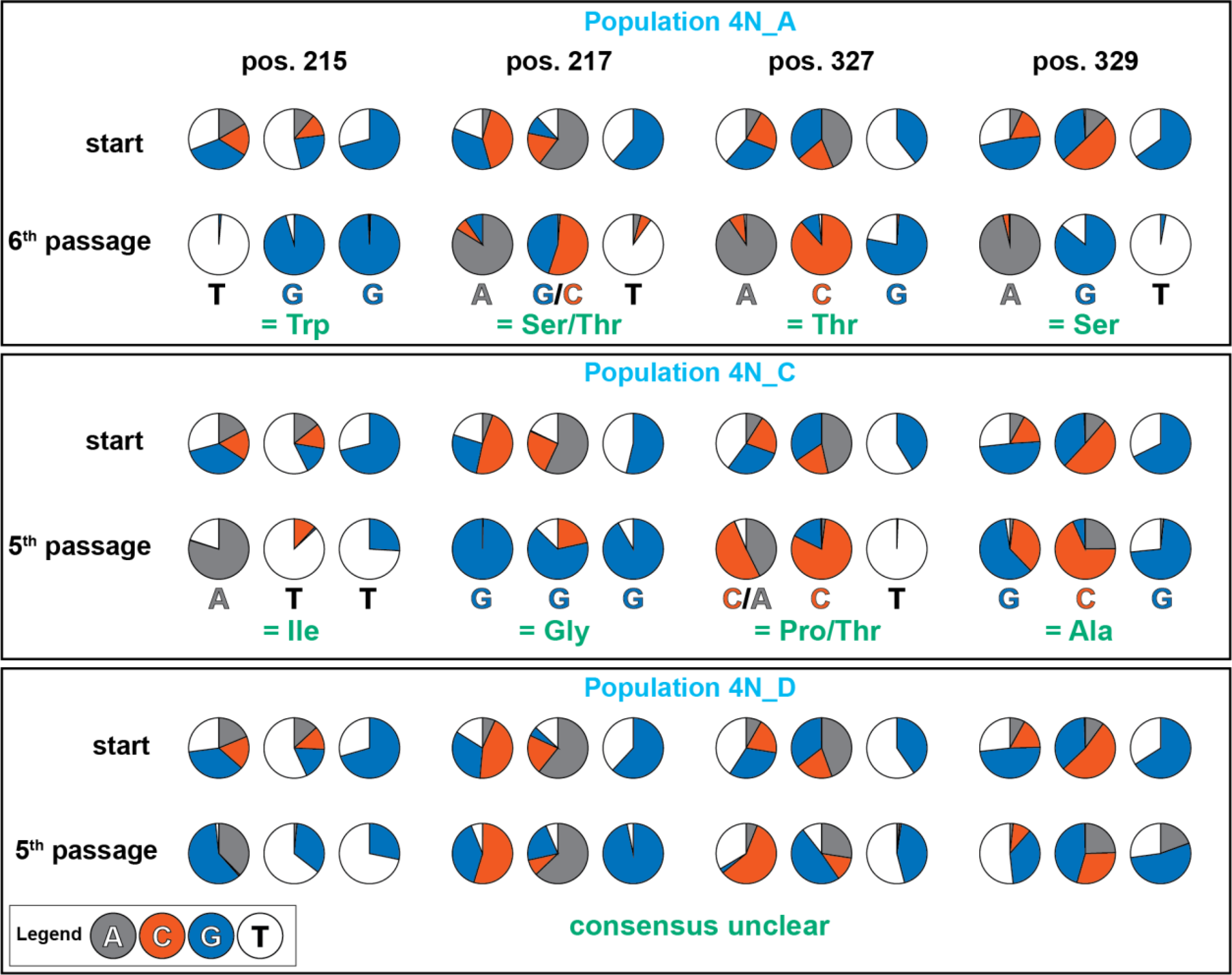
Genotypical changes after selection. Pie charts depict the genotypes of populations 4N_A, 4N_C, and 4N_D after 5 or 6 serial passages derived from the base calls obtained from Sanger sequencing pooled plasmids. The consensus of the genotypes is given in green.

**Extended Data Fig. 5:**
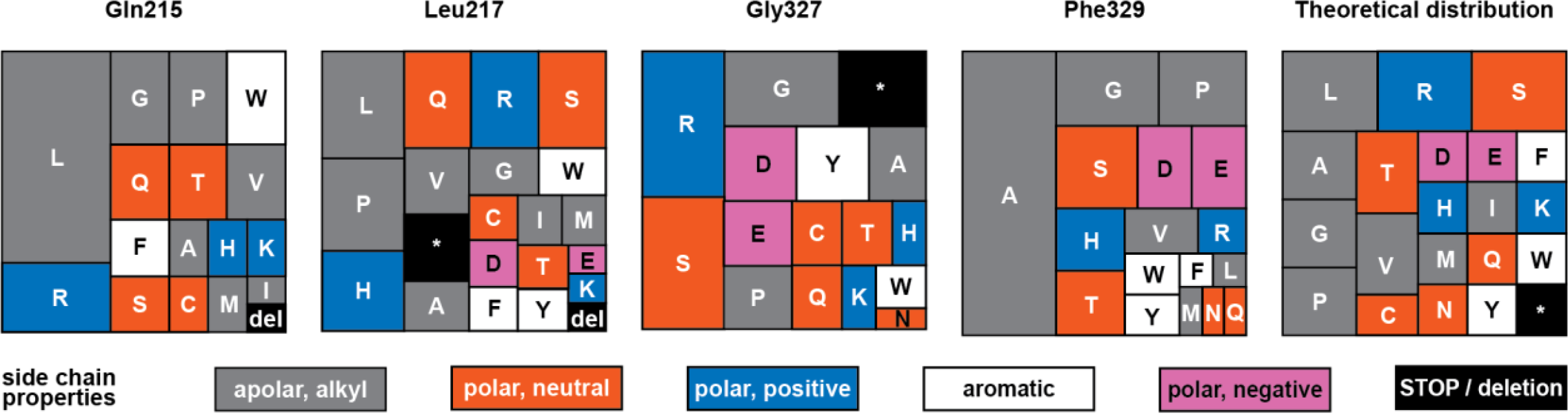
Amino acid distribution of Lib_4N before selection. Tree map charts depicting the amino acid distribution across all four randomized positions in Lib_4N based on sequencing data of 73 single colonies prior to selection. The theoretical distribution is based on the perfect NNK codon. See also Supplementary Table S6.

**Extended Data Fig. 6:**
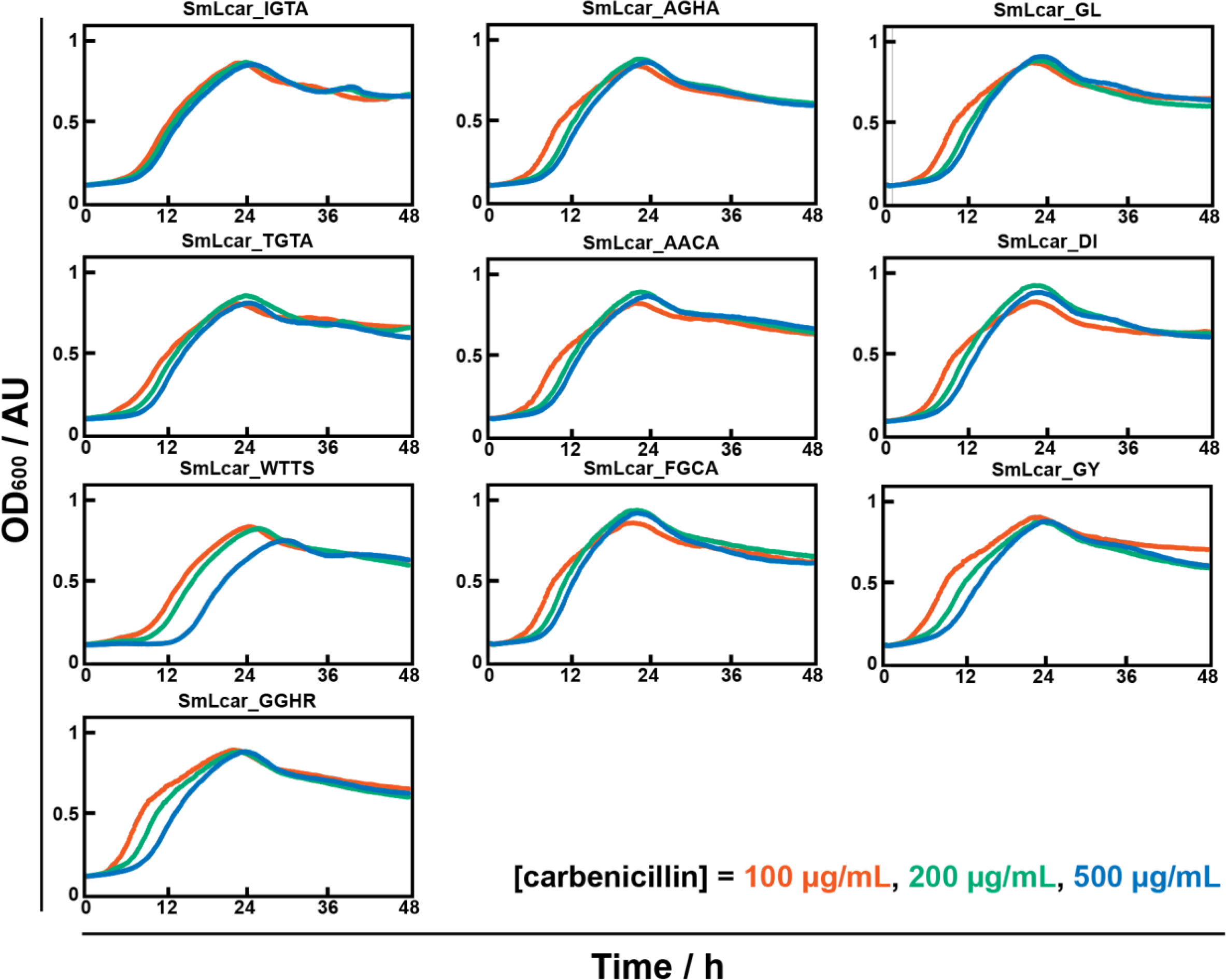
Growth curves of selected SmLcar variants that originated from Lib_2N and Lib_4N at increasing carbenicillin concentrations. SmLcar_GY is included as a reference for high carbamoylase activity. SmLcar variants of interest were cloned and transformed into new recoded *E. coli* hosts. Note that the lag time increases slightly with increasing carbenicillin pressure. Cells were grown in the presence of 500 μM cam-3nY.

