## Supplementary Information for "A robust life-or-death selection platform for enzyme evolution"

**Affiliations:**

#### Table of contents

1. Supplementary Figures S2
2. Supplementary Tables S4
3. Methods S13
4. Sequences S26
5. Supplementary References S32

### 1. Supplementary Figures

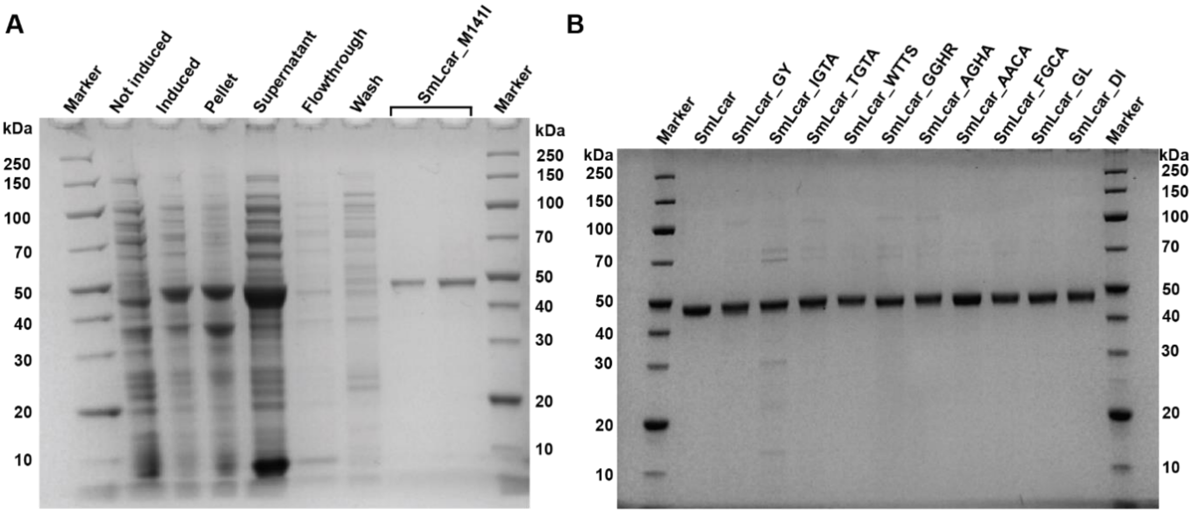


**Supplementary Fig. S1: SDS-PAGE analysis of purified SmLcar variants. A:** SDS-PAGE gel depicting the successful purification of SmLcar_M141I (~47 kDa). Purifications of other SmLcar variants look similar, so for clarity, only one mutant is shown. **B:** Elution fractions of selected SmLcar variants (~47 kDa) show that the enzymes were obtained in high purity.

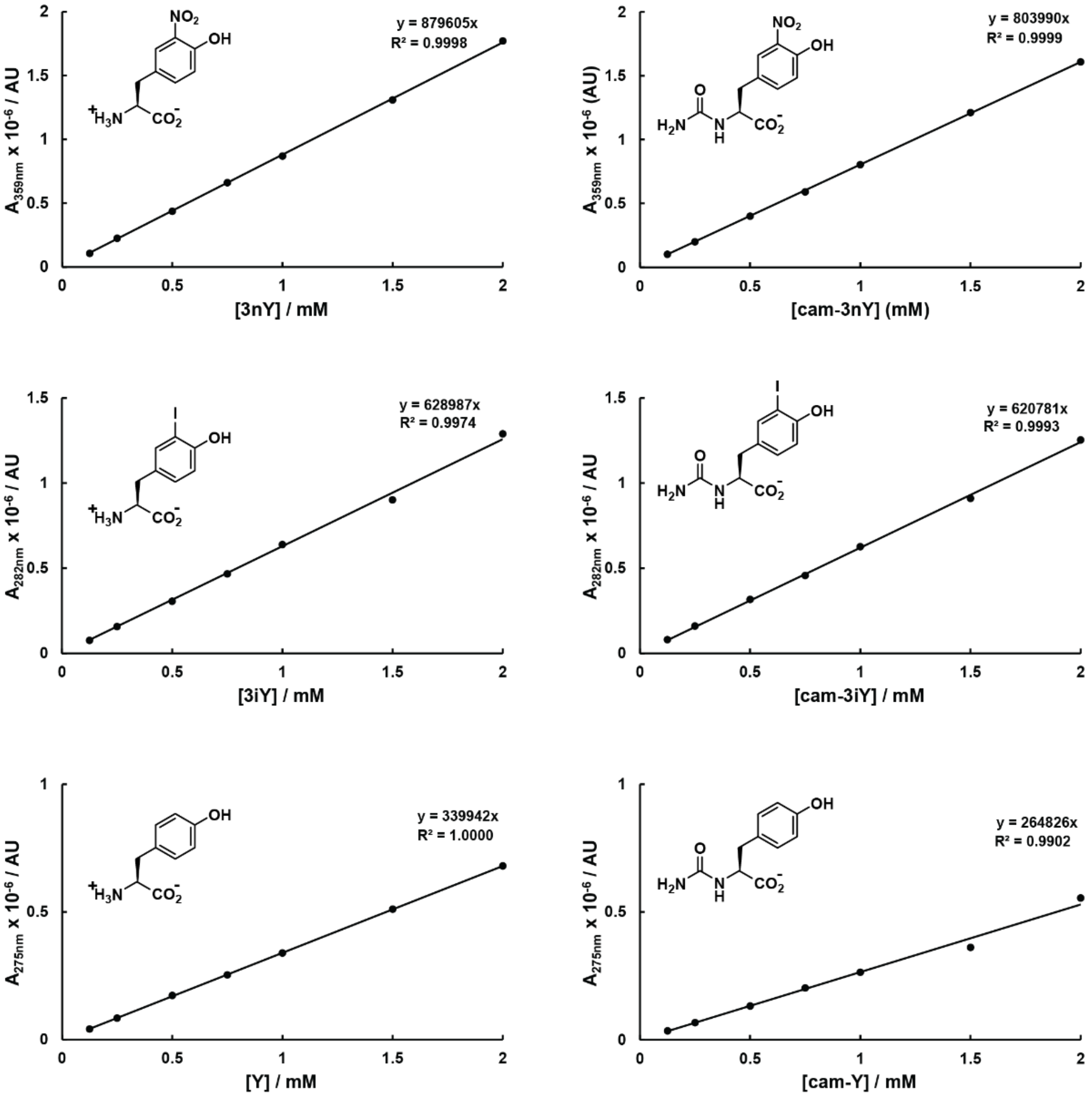


**Supplementary Fig. S2: HPLC calibration curves for (nc)AAs and carbamoylated precursors.** Chemical structures, trendlines and $R^2$-values are given in the graphs.

### 2. Supplementary Tables

**Supplementary Table S1: Transformation efficiencies.** The transformation efficiency was calculated based on the number of colony-forming units and final culture volume for each library.

| | Host | Method | µg DNA added | cfu on diluted plate | Dilution | # transformants |
|---|---|---|---|---|---|---|
| **Lib_2N** | NEB10-beta | Chemical transformation | 0.1 | 2432 | None.<br>3900 µL was concentrated and plated.<br>Total cell volume: 3900 µL | **2432** |
| **Lib_4N** | NEB10-beta | Electroporation | 0.306 | 40 | $10^{-3}$ dilution.<br>180 µl was plated.<br>Total cell volume: 3800 µL. | 40 cfu / 180 µL * 200 µL * $10^3$ / 20 µL * 3800 µL = **8.4 *$10^6$** |
| **Pop. 4N_A** | BL21(DE3) | Electroporation | 0.3 | 516 | $10^{-3}$ dilution.<br>180 µl was plated.<br>Total cell volume: 2000 µL | 516 cfu / 180 µL * 200 µL * $10^3$ / 20 µL * 2000 µL = **5.7 * $10^7$** |
| **Pop. 4N_B** | BL21(DE3) | Electroporation | 0.3 | 388 | $10^{-3}$ dilution.<br>180 µl was plated.<br>Total cell volume: 2000 µL | 388 cfu / 180 µL * 200 µL * $10^3$ / 20 µL * 2000 µL = **4.3 * $10^7$** |
| **Pop. 4N_C** | BL21(DE3) | Electroporation | 0.3 | 137 | $10^{-3}$ dilution.<br>180 µl was plated.<br>Total cell volume: 3800 µL | 137 cfu / 180 µL * 200 µL * $10^3$ / 20 µL * 3800 µL = **2.9 * $10^7$** |
| **Pop. 4N_D** | BL21(DE3) | Electroporation | 0.3 | 169 | $10^{-3}$ dilution.<br>180 µl was plated.<br>Total cell volume: 3800 µL | 169 cfu / 180 µL * 200 µL * $10^3$ / 20 µL * 3800 µL = **3.6 * $10^7$** |

**Supplementary Table S2: Library quality.** Base read frequencies and resulting Q-values[1] for each randomized NNK stretch in starting populations, as followed from Sanger sequencing. The base read frequencies are based on pooled plasmids for Lib_2N and aggregate data of 71 single colonies for Lib_4N. Note that a perfect NNK – with both Ns containing 25% of each base, and the K containing 50% G and 50% T – would give a Q-value of 1.

| Library | Randomized codon | Base read frequency (%) | | | | Q-value |
|---|---|---|---|---|---|---|
| | | A | C | G | T | |
| Lib_2N | L217_**N**NK | 16.8 | 36.9 | 26.4 | 19.9 | 0.85 |
| | L217_N**N**K | 21.7 | 22.7 | 19.2 | 36.5 | |
| | L217_NN**K** | 0 | 0 | 47.5 | 52.5 | |
| | F329_**N**NK | 14.3 | 40.6 | 17.6 | 27.5 | 0.82 |
| | F329_N**N**K | 35.5 | 15.4 | 24.4 | 24.7 | |
| | F329_NN**K** | 0 | 0 | 43.6 | 56.4 | |
| Lib_4N | Q215_**N**NK | 14.1 | 42.3 | 15.5 | 28.2 | 0.74 |
| | Q215_N**N**K | 9.9 | 19.7 | 25.4 | 45.1 | |
| | Q215_NN**K** | 0 | 0 | 57.7 | 42.3 | |
| | L217_**N**NK | 11.3 | 43.7 | 18.3 | 26.8 | 0.82 |
| | L217_N**N**K | 32.4 | 23.9 | 18.3 | 25.4 | |
| | L217_NN**K** | 0 | 0 | 42.3 | 57.7 | |
| | G327_**N**NK | 18.3 | 23.9 | 28.2 | 29.6 | 0.80 |
| | G327_N**N**K | 40.8 | 23.9 | 35.2 | 0 | |
| | G327_NN**K** | 1.4 | 0 | 53.5 | 45.1 | |
| | F329_**N**NK | 9.9 | 16.9 | 57.7 | 15.5 | 0.63 |
| | F329_N**N**K | 22.5 | 54.9 | 14.1 | 8.5 | |
| | F329_NN**K** | 0 | 0 | 54.9 | 45.1 | |

**Supplementary Table S3: Random mutations.** Random copying errors found in 71 individual clones from Lib_4N, outside of the targeted NNK codons. Of the 53,037 bases analyzed, 30 had undergone a random mutation. Detailed analysis is available in Supplementary Data File 1.

| **Nature of random mutation** | | **# found** | **Share of total random mutations (%)** |
|---|---|---|---|
| Correctly replicated | | 53,007 | N/A |
| Transition | A ⇄ G | 15 | 50 |
| | C ⇄ T | 8 | 27 |
| Transversion | G ⇄ T | 3 | 10 |
| | C ⇄ A | 3 | 10 |
| | T ⇄ A | 1 | 3 |
| | G ⇄ C | 0 | 0 |

**Supplementary Table S4: Selecting for growth advantage.** Growth characteristics found in individual library members taken from the start and endpoints of several evolutionary trajectories. No growth is defined as failing to exceed an $OD_{600}$ of 0.3 within 48 hours; slow growth is defined as exceeding an $OD_{600}$ of 0.3 after 24 to 48 hours; fast growth is defined as exceeding an $OD_{600}$ of 0.3 within 24 hours.

| Population | Carbenicillin concentration (µg/mL) | Total # of colonies | Growth characteristics (counts, percentages) | | | | | |
|---|---|---|---|---|---|---|---|---|
| | | | No growth | | Slow growth | | Fast growth | |
| 2N Gen 0 | 100 | 95 | 54 | 57% | 23 | 24% | 18 | 19% |
| | 500 | | 82 | 86% | 11 | 12% | 2 | 2% |
| 2N_A Gen 3 | 100 | 47 | 4 | 9% | 4 | 9% | 39 | 83% |
| | 500 | | 9 | 19% | 5 | 11% | 33 | 70% |
| 2N_B Gen 4 | 100 | 48 | 5 | 10% | 1 | 2% | 42 | 88% |
| | 500 | | 9 | 19% | 0 | 0% | 39 | 81% |
| 4N Gen 0 | 100 | 95 | 85 | 89% | 7 | 7% | 3 | 3% |
| | 500 | | 95 | 100% | 0 | 0% | 0 | 0% |
| 4N_A1 Gen 6 | 100 | 95 | 6 | 6% | 0 | 0% | 89 | 94% |
| | 500 | 127 | 43 | 34% | 58 | 46% | 26 | 20% |
| 4N_A2 Gen 5 | 100 | 95 | 70 | 74% | 1 | 1% | 24 | 25% |
| | 500 | | 77 | 81% | 17 | 18% | 1 | 1% |
| 4N_C Gen 5 | 100 | 95 | 1 | 1% | 4 | 4% | 90 | 95% |
| | 500 | 127 | 14 | 11% | 4 | 3% | 109 | 86% |
| 4N_D1 Gen 5 | 100 | n.d. | n.d. | n.d. | n.d. | n.d. | n.d. | n.d. |
| | 500 | 31 | 17 | 55% | 1 | 3% | 13 | 42% |
| 4N_D2 Gen 5 | 100 | 95 | 0 | 0% | 10 | 11% | 85 | 89% |
| | 500 | | 12 | 13% | 15 | 16% | 68 | 72% |

**Supplementary Table S5: Sequencing results of the best-performing colonies of population 2N_A (Gen 3) and 2N_B (Gen 4).** For 2N_A, clones 9 and 10 are removed due to unfaithful sequencing traces. Variants chosen for characterization are marked in green. For the additional mutations and substitutions, deviations from the wildtype sequence are indicated in bold.

| Population | Clone no. | Phenotype shorthand | codon 217 | L217 | codon 329 | F329 | Additional substitutions and mutations |
|---|---|---|---|---|---|---|---|
| 2N_A Gen 3 | 1 | GY | GGT | Gly | TAT | Tyr | none |
| | 2 | GY | GGT | Gly | TAT | Tyr | none |
| | 3 | GY | GGT | Gly | TAT | Tyr | silent mutation A310A (GC**G**→GC**A**) |
| | 4 | GY | GGT | Gly | TAT | Tyr | mutation S233T (**T**CA→**A**CA) |
| | 5 | GY | GGT | Gly | TAT | Tyr | silent mutation A310A (GC**G**→GC**A**) |
| | 6 | GY | GGT | Gly | TAT | Tyr | none |
| | 7 | GY | GGT | Gly | TAT | Tyr | silent mutation S86S (TC**G**→TC**A**), mutation E221D (GA**A**→GA**T**), mutation A346T (**G**CA→**A**CA) |
| | 8 | GY | GGT | Gly | TAT | Tyr | silent mutation S86S (TC**G**→TC**A**), mutation A346T (**G**CA→**A**CA) |
| 2N_B Gen 4 | 1 | DI | GAT | Asp | ATT | Ile | mutation M141I (AT**G**→AT**A**), mutation Q296R (C**A**A→C**G**A), silent mutation A401A (GC**A**→GC**G**) |
| | 2 | GY | GGT | Gly | TAT | Tyr | none |
| | 3 | DI | GAT | Asp | ATT | Ile | mutation M141I (AT**G**→AT**A**), mutation Q296R (C**A**A→C**G**A), silent mutation A401A (GC**A**→GC**G**) |
| | 4 | GL | GGG | Gly | CTT | Leu | mutation M141I (AT**G**→AT**A**) |
| | 5 | DI | GAT | Asp | ATT | Ile | mutation M141I (AT**G**→AT**A**), mutation Q296R (C**A**A→C**G**A), silent mutation A401A (GC**A**→GC**G**) |
| | 6 | DI | GAT | Asp | ATT | Ile | mutation M141I (AT**G**→AT**A**), mutation Q296R (C**A**A→C**G**A), silent mutation A401A (GC**A**→GC**G**) |
| | 7 | DI | GAT | Asp | ATT | Ile | mutation M141I (AT**G**→AT**A**), mutation Q296R (C**A**A→C**G**A), silent mutation A401A (GC**A**→GC**G**) |
| | 8 | GL | GGG | Gly | CTT | Leu | mutation M141I (AT**G**→AT**A**) |
| | 9 | DI | GAT | Asp | ATT | Ile | mutation M141I (AT**G**→AT**A**), mutation Q296R (C**A**A→C**G**A), silent mutation A401A (GC**A**→GC**G**) |
| | 10 | GL | GGG | Gly | CTT | Leu | mutation M141I (AT**G**→AT**A**) |

**Supplementary Table S6: Sequencing results and growth characteristics of 4N library members prior to selection.** Sequencing results of 73 single colonies taken from 4N_A, 4N_C and 4N_D populations, prior to selection based on growth advantage. Table continues on next page.

| entry | Growth category (at 500 ug/ml carb) | Source population | 4-letter | Q215 | L217 | G327 | F329 | additional substi-tutions | # found |
|---|---|---|---|---|---|---|---|---|---|
| 1 | no growth | A Gen0 | AIDA | A | I | D | A | none | 1 |
| 2 | no growth | A Gen0 | ARDD | A | R | D | D | M238I | 1 |
| 3 | no growth | A Gen0 | CP*H | C | P | * | H | none | 1 |
| 4 | no growth | D Gen0 | CQKP | C | Q | K | P | none | 1 |
| 5 | no growth | A Gen0 | --CW | - | - | C | W | residues 215, 216, and 217 are deleted | 1 |
| 6 | no growth | D Gen0 | FHGA | F | H | G | A | H328N | 1 |
| 7 | no growth | C Gen0 | FRSG | F | R | S | G | none | 1 |
| 8 | no growth | C Gen0 | F**A | F | * | * | A | G232R | 1 |
| 9 | no growth | C Gen0 | GF*E | G | F | * | E | R227H | 1 |
| 10 | no growth | D Gen0 | GGTY | G | G | T | Y | none | 1 |
| 11 | no growth | D Gen0 | GHRA | G | H | R | A | A304T | 1 |
| 12 | no growth | D Gen0 | GPWR | G | P | W | R | none | 1 |
| 13 | no growth | D Gen0 | G*EH | G | * | E | H | R227H | 1 |
| 14 | no growth | D Gen0 | HCYQ | H | C | Y | Q | none | 1 |
| 15 | no growth | C Gen0 | HMGA | H | M | G | A | none | 1 |
| 16 | no growth | A Gen0 | IREE | I | R | E | E | none | 1 |
| 17 | no growth | C Gen0 | KSEA | K | S | E | A | none | 1 |
| 18 | no growth | C Gen0 | KSQP | K | S | Q | P | none | 1 |
| 19 | no growth | C Gen0 | LAEP | L | A | E | P | none | 1 |
| 20 | no growth | D Gen0 | LASA | L | A | S | A | none | 1 |
| 21 | no growth | A Gen0 | LDAY | L | D | A | Y | none | 1 |
| 22 | no growth | D Gen0 | LHGS | L | H | G | S | P162Q | 1 |
| 23 | no growth | D Gen0 | LIRA | L | I | R | A | none | 1 |
| 24 | no growth | A Gen0 | LLDA | L | L | D | A | none | 1 |
| 25 | no growth | D Gen0 | LLSF | L | L | S | F | G198D | 1 |
| 26 | no growth | C Gen0 | LLSW | L | L | S | W | none | 1 |
| 27 | no growth | C Gen0 | LPKG | L | P | K | G | R305C | 1 |
| 28 | no growth | A Gen0 | LPSR | L | P | S | R | none | 1 |
| 29 | no growth | A Gen0 | LPTN | L | P | T | N | V268M | 1 |
| 30 | no growth | A Gen0 | LP*P | L | P | * | P | none | 1 |
| 31 | no growth | A Gen0 | LQND | L | Q | N | D | none | 1 |
| 32 | no growth | C Gen0 | LQYP | L | Q | Y | P | A247S | 1 |
| 33 | no growth | D Gen0 | LSQA | L | S | Q | A | none | 1 |
| 34 | no growth | C Gen0 | LVQA | L | V | Q | A | none | 1 |
| 35 | no growth | C Gen0 | LVSA | L | V | S | A | none | 1 |
| 36 | no growth | D Gen0 | LWAA | L | W | A | A | none | 1 |
| 37 | no growth | A Gen0 | LWRV | L | W | R | V | none | 1 |
| 38 | no growth | D Gen0 | LYPE | L | Y | P | E | none | 1 |
| 39 | no growth | C Gen0 | LYYA | L | Y | Y | A | S352L | 1 |

**Supplementary Table S6:** continued.

| entry | Growth category (at 500 ug/ml carb) | Source population | 4-letter | Q215 | L217 | G327 | F329 | additional substi-tutions | # found |
|---|---|---|---|---|---|---|---|---|---|
| 40 | no growth | A Gen0 | MQSV | M | Q | S | V | A360T | 1 |
| 41 | no growth | A Gen0 | MTWD | M | T | W | D | none | 1 |
| 42 | no growth | A Gen0 | PHPA | P | H | P | A | none | 1 |
| 43 | no growth | A Gen0 | PLRS | P | L | R | S | none | 1 |
| 44 | no growth | C Gen0 | PQHE | P | Q | H | E | M258I | 1 |
| 45 | no growth | C Gen0 | PVYG | P | V | Y | G | none | 1 |
| 46 | no growth | C Gen0 | P*CM | P | * | C | M | none | 1 |
| 47 | no growth | A Gen0 | QCCA | Q | C | C | A | none | 1 |
| 48 | no growth | D Gen0 | QGGA | Q | G | G | A | none | 1 |
| 49 | no growth | D Gen0 | QLAL | Q | L | A | L | none | 1 |
| 50 | no growth | D Gen0 | QRGA | Q | R | G | A | G216D | 1 |
| 51 | no growth | D Gen0 | RAHG | R | A | H | G | none | 1 |
| 52 | no growth | A Gen0 | RFRT | R | F | R | T | none | 1 |
| 53 | no growth | A Gen0 | RKDG | R | K | D | G | none | 1 |
| 54 | no growth | A Gen0 | RMTA | R | M | T | A | none | 1 |
| 55 | no growth | A Gen0 | RPPT | R | P | P | T | Q296H | 1 |
| 56 | no growth | D Gen0 | RRPH | R | R | P | H | A148V | 1 |
| 57 | no growth | D Gen0 | RSSS | R | S | S | S | none | 1 |
| 58 | no growth | A Gen0 | SHSG | S | H | S | G | none | 1 |
| 59 | no growth | C Gen0 | SLRP | S | L | R | P | R248H | 1 |
| 60 | no growth | C Gen0 | SQ*A | S | Q | * | A | none | 1 |
| 61 | no growth | C Gen0 | TD*G | T | D | * | G | none | 1 |
| 62 | no growth | C Gen0 | TSDD | T | S | D | D | S322P | 1 |
| 63 | no growth | D Gen0 | TVYA | T | V | Y | A | none | 1 |
| 64 | no growth | D Gen0 | T*RS | T | * | R | S | none | 1 |
| 65 | no growth | D Gen0 | VGST | V | G | S | T | A360V | 1 |
| 66 | no growth | C Gen0 | VLRS | V | L | R | S | A346T & W218* | 1 |
| 67 | no growth | C Gen0 | VRGS | V | R | G | S | none | 1 |
| 68 | no growth | D Gen0 | VTAH | V | T | A | H | none | 1 |
| 69 | no growth | D Gen0 | WEGV | W | E | G | V | none | 1 |
| 70 | no growth | A Gen0 | WHGA | W | H | G | A | none | 1 |
| 71 | no growth | D Gen0 | WLRA | W | L | R | A | none | 1 |
| 72 | no growth | D Gen0 | WSRA | W | S | R | A | none | 1 |
| 73 | no growth | C Gen0 | WWRT | W | W | R | T | Y351D | 1 |

**Supplementary Table S7: Selection has enriched 37 fast-growing variants.** Sequencing results of 132 single colonies taken from 4N_A, 4N_C and 4N_D populations after selection reveal 37 unique variants. Variants chosen for characterization are marked in green.

| entry | Growth category (at 500 ug/ml carb) | Source population | 4-letter | Q215 | L217 | G327 | F329 | additional substi-tutions | # found |
|---|---|---|---|---|---|---|---|---|---|
| 1 | Fast | C Gen5 | AACA | A | A | C | A | none | 2 |
| 2 | Fast | A1 Gen6 & C Gen5 | FACA | F | A | C | A | R303C | 2 |
| 3 | Fast | D1 Gen5 | FACA | F | A | C | A | none | 1 |
| 4 | Fast | C Gen5 | FACS | F | A | C | S | none | 2 |
| 5 | Fast | C Gen5 | GACN | G | A | C | N | none | 1 |
| 6 | Fast | D1 Gen5 | IACC | I | A | C | C | H353Y | 3 |
| 7 | Fast | C Gen5 | IAHR | I | A | H | R | none | 1 |
| 8 | Fast | D1 Gen5 | LACT | L | A | C | T | none | 1 |
| 9 | Fast | C Gen5 | SACA | S | A | C | A | none | 1 |
| 10 | Fast | C Gen5 | SAGA | S | A | G | A | none | 2 |
| 11 | Fast | A1 Gen6 & A2 Gen5 | TACA | T | A | C | A | none | 2 |
| 12 | Fast | C Gen5 | TACC | T | A | C | C | none | 1 |
| 13 | Fast | C Gen5 | TAHR | T | A | H | R | A368T & V326A | 1 |
| 14 | Fast | A1 Gen6 | TASG | T | A | S | G | E180G & A138V | 1 |
| 15 | Fast | A1 Gen6 & C Gen5 | VACA | V | A | C | A | none | 2 |
| 16 | Fast | D1 Gen5 | VACC | V | A | C | C | none | 1 |
| 17 | Fast | D1 Gen5 | VACS | V | A | C | S | none | 1 |
| 18 | Fast | A1 Gen6 & A2 Gen5 | VAHR | V | A | H | R | none | 2 |
| 19 | Fast | D1 Gen5 | WATA | W | A | T | A | V326A | 1 |
| 20 | Fast | C Gen5 | AGHA | A | G | H | A | A368T | 1 |
| 21 | Fast | A2 Gen5 | FGCA | F | G | C | A | M141I | 1 |
| 22 | Fast | A1 Gen6 | FGCV | F | G | C | V | A367V | 1 |
| 23 | Fast | C Gen5 | GGHR | G | G | H | R | I208M & I200F | 1 |
| 24 | Fast | C Gen5 | GGHR | G | G | H | R | F191S | 1 |
| 25 | Fast | C Gen5 | GGHR | G | G | H | R | none | 2 |
| 26 | Fast | C Gen5 | GGTA | G | G | T | A | E315G & G175D | 1 |
| 27 | Fast | C Gen5 | GGTR | G | G | T | R | none | 1 |
| 28 | Fast | C Gen5 | IGTA | I | G | T | A | A368T | 40 |
| 29 | Fast | D1 Gen5 | QGHR | Q | G | H | R | none | 1 |
| 30 | Fast | A2 Gen5 | SGCV | S | G | C | V | P262S | 1 |
| 31 | Fast | D1 Gen5 | TGHR | T | G | H | R | none | 1 |
| 32 | Fast | D2 Gen5 | TGTA | T | G | T | A | none | 30 |
| 33 | Fast | C Gen5 | VGHR | V | G | H | R | A346T | 1 |
| 34 | Fast | D1 Gen5 | LSCA | L | S | C | A | M141I | 1 |
| 35 | Fast | D1 Gen5 | WSGR | W | S | G | R | none | 1 |
| 36 | Fast | A1 Gen6 | WTTS | W | T | T | S | none | 18 |
| 37 | Fast | A1 Gen6 | WTTS | W | T | T | S | A143P | 1 |

**Supplementary Table S8: Mass analysis of purified enzymes.** Q-TOF MS results confirm the identity of several purified SmLcar variants.

| Carbamoylase variant | expected (M-Met, Da) | observed (M-Met, Da) |
|---|---|---|
| **SmLcar wildtype** | 46850.13 | 46849 |
| **SmLcar_GY** | 46840.05 | 46839 |
| **SmLcar_M141I** | 46832.10 | 46833 |
| **SmLcar_DI** | 46828.07 | 46828 |
| **SmLcar_GL** | 46741.97 | 46741 |
| **SmLcar_AACA** | 46720.99 | 46722 |
| **SmLcar_FGCA** | 46765.02 | 46764 |
| **SmLcar_IGTA** | 46777.03 | 46776 |
| **SmLcar_GGHR** | 46812.04 | 46809 |
| **SmLcar_TGTA** | 46734.95 | 46733 |
| **SmLcar_AGHA** | 46770.99 | 46769 |
| **SmLcar_WTTS** | 46880.11 | 46879 |

#### 3. Methods

**Materials & Methods:** Chemicals, including L-3-nitro-tyrosine (3nY), L-3-iodo-tyrosine (3iY), L-tyrosine (Tyr) and cam-L-tyrosine (cam-Y), were purchased from *Sigma Aldrich* and used without further purification unless noted otherwise. *Escherichia coli* strains NEB 10-beta and BL21(DE3) (*New England Biolabs*) were used for cloning and expression experiments, respectively. Bacteria were cultured in Lysogeny broth (LB) medium, or Super Optimal Broth (with Catabolite repression) (SOB, SOC) medium during bacterial cell transformation. Standard and sequencing primers were synthesized by *Eurofins Genomics* (Germany), while primers with degenerate codons were synthesized by *Biolegio* (The Netherlands). Plasmid isolation kits (QIAprep Spin Miniprep Kits) and PCR and gel clean-up kits (QIAquick PCR Purification Kit, QIAquick Gel Extraction Kit) were purchased from *QIAGEN* (Germany). Sanger sequencing of plasmids and PCR products was carried out by *Eurofins Genomics* (Germany) using TubeSeq NXP Service. Phusion® High Fidelity DNA Polymerase, GC buffer, deoxynucleotide (dNTP) solution mix, Hi-T4 DNA ligase, T4 DNA ligase reaction buffer, dimethyl sulfoxide (DMSO), *BsaI*-HF, *EcoRI*-HF, *XhoI*, *DpnI* and CutSmart® buffer were purchased from *New England Biolabs*. GeneRuler 1kb DNA ladder, TriTrack DNA loading dye, Orange DNA loading dye, and PageRuler™ Unstained Broad Range Protein Ladder were purchased from *ThermoFisher Scientific*. ROTI®GelStain was purchased from *Carl Roth* (Germany). Ni-NTA resin (Ni Sepharose 6 Fast Flow) was purchased from *Cytiva* (Germany). Precast SDS PAGE gels (ExpressPlus™ PAGE Gel, 10x8, 12%, 15 wells) and Tris-MOPS-SDS Running Buffer Powder were purchased from *GenScript* (USA). Electroporation cuvettes (2 mm) were purchased from *Fisher Scientific* (The Netherlands).

The concentration of DNA in solutions was determined based on the absorption at 260 nm on a Thermo Scientific Nanodrop 2000 UV-Vis spectrophotometer. The concentration of protein in solutions was determined based on the absorption at 280 nm on the same apparatus.

Cellular density ($OD_{600}$) was measured on an Ultrospec 10 Cell Density Meter (Biochrom). Plate reader growth assays were recorded on a Synergy H1 microplate reader (*BioTek*). Agarose or SDS-PAGE gels were imaged with a GelDoc Go Imaging System (Biorad) equipped with an UV tray or white tray, respectively. Analytical HPLC analysis was performed on a Waters Acquity HPLC class system (Waters) equipped with a PDA detector. All analyses were performed using a reverse-phase HPLC column (XSelect-CSH-C18, 5 µm, 4.5×150 mm; Waters) kept at 40 °C and the sample plate was kept at room temperature. Samples were separated with a gradient from 5 to 95% acetonitrile (0.1% TFA) in Milli-Q water (0.1% TFA) at a flow rate of 1.0 mL/min, each run lasting 27 minutes. Absorbance was monitored at different wavelengths for the different (nc)AAs and their carbamoylated derivatives (359 nm for 3nY, 282 nm for 3iY, 275 nm for Tyr). Each (nc)AA or cam-(nc)AA was quantified according to a calibration curve obtained from samples containing different concentrations of authentic standards (**Supplementary Fig. S2**).

**Synthesis of cam-3iY and cam-3nY**: Synthesis of the carbamoylated ncAA precursors, cam-L-3-iodo-tyrosine (cam-3iY) and cam-L-3-nitro-tyrosine (cam-3nY), and characterization of the compounds via $^{1}$H-NMR and $^{13}$C-NMR has been described in our previous work.[2]

**Construction of pULTRA_3iY and pACYC_GG:** Construction of the OTS plasmid pULTRA_3iY and the selection plasmid pACYC_GG, based on the commercially available pACYCDuet-1, has been described in our previous work.[2] In brief, pULTRA-3iY is based on pULTRA-CNF[3], but the CNF aminoacyl tRNA synthetase (aaRS) is exchanged for that of the 3iY-aaRS while retaining the same tRNA cassette (see *Sequences*). In the dual expression vector pACYC_GG, modular exchange of the target enzyme and/or the beta-lactamase in both multiple cloning sites (MCSs) has been constructed. PCR products and/or synthetic genes flanked by the appropriate type II2 recognition sites can be easily assembled into either MCS1 (using *BsaI*) or MCS2 (using *Esp3I*) of pACYC_GG with Golden Gate Assembly. The current work uses pACYC_GG (with two available MCSs) or pACYC_TEM-1.B9 (in which MCS2 is occupied by the CDS of TEM-1.B9). In all cases, the target enzymes, i.e. SmLcar variants and libraries, were cloned into MCS1. The SmLcar CDS and TEM-1.B9 CDS were already in our possession from previous work where they were cloned into pACYC_GG (see *Sequences*).[2]

In order to realize the chemical complementation system, a selection plasmid was co-transformed (either chemically or through electroporation) in *E. coli* BL21(DE3) already bearing the OTS plasmid pULTRA_3iY (*vide infra*).

**Site-directed mutagenesis:** Starting from selection plasmids pACYC_SmLcar_Leu217Gly-Phe329Tyr (pACYC_SmLcar_GY) and pACYC_SmLcar_TEM-1.B9, mutagenic primers were used to generate digestive selection plasmids pACYC_SmLcar_GY* and pACYC_SmLcar** with one and two restriction sites, respectively. First, mutagenic primers *SmLcar_TTT-TTC_F222F_fw* and *SmLcar_TTT-TTC_F222F_rv* were used to create an *EcoRI* restriction site in the SmLcar CDS of both templates by introducing a silent mutation (F222F). The following PCR protocol was used for the pACYC_SmLcar_GY template: (1) initial denaturation at 95 ˚C for 3 min, (2) 16 cycles of denaturation at 95 ˚C for 30 s, annealing at 58 ˚C for 30 s, and extension at 72 ˚C for 3 min; (3) a final extension at 72 ˚C for 10 min. For the pACYC_SmLcar_TEM-1.B9 template, the same PCR protocol was used except the annealing temperature was 61˚C. The resulting PCR products were digested with *DpnI* for 1 hour at 37 ˚C, purified, and transformed into chemically competent *E. coli* NEB10-beta cells. A single colony was picked from LB plates containing chloramphenicol (35 µg/mL) and was used to inoculate 4 mL of LB medium containing the same concentration of chloramphenicol. Bacteria were grown overnight, plasmids were isolated and variants harboring the correct mutations were identified by sequencing with *DuetDOWN1*. The resulting plasmids are pACYC_SmLcar_F222F and pACYC_SmLcarF222F_GY (=pACYC_SmLcarGY*_TEM-1.B9). The former plasmid was subjected to additional site-directed mutagenesis.

To introduce the second restriction site, pACYC_SmLcar_F222F_TEM-1.B9 was used as a template. A *XhoI* site was introduced into the non-coding region of the vector between the p15A origin of replication and lacI CDS using mutagenic primers *SmLcar_GTC-CTC_QC_fw* and *SmLcar_GTC-CTC_QC_rv*. The same PCR protocol was used as above, now with an annealing temperature of 64 ˚C. *DpnI* digestion, PCR cleanup, transformation, and overnight growth was performed as described above. Variants harboring the correct mutations were identified by sequencing with *pACYC_GG_3rev*. The resulting plasmid is shortened for convenience and named pACYC_SmLcar**.

pACYC_SmLcar** and pACYC_SmLcar_GY* both have TEM-1.B9 at the second MCS of pACYC, and these selection plasmids were transformed into chemically competent *E. coli* BL21(DE3) cells harboring pULTRA_3iY and used to perform mock selections.

To construct a variant of SmLcar with the Met141Ile substitution, plasmids pACYC_SmLcar and pACYC_SmLcar_TEM-1.B9 were used as a template with mutagenic primers *SmLcar_M141I_fw* and *SmLcar_M141_rev*. The same PCR protocol as above was used, now with an annealing temperature of 65 ˚C. *DpnI* digestion, PCR cleanup, transformation, and overnight growth was performed as described above. Variants harboring the correct mutation were identified by sequencing with *DuetDOWN1*. pACYC_SmLcarM141 was chemically transformed into *E. coli* BL21(DE3), and used for producing the enzyme variant for purification. pACYC_SmLcarM141I_TEM-1.B9 was chemically transformed into *E. coli* BL21(DE3) already bearing the OTS plasmid pULTRA_3iY and chemical complementation was tested in 96-well plates (*vide infra*).

**Mock selections:** Precultures of 5 mL LB media containing 50 µg/mL spectinomycin and 35 µg/ml chloramphenicol were inoculated from fresh agar plates with *E. coli* BL21(DE3) cells harboring pULTRA-3iY and the selection plasmid featuring either SmLcar** or SmLcar_GY*. Following overnight growth at 37 °C while shaking at 135 rpm, a main culture of 5 mL LB containing 25 µg/ml spectinomycin and 17.5 µg/ml chloramphenicol was started with SmLcar** cells and SmLcar_GY* cells mixed in a 100:1 ratio (specifically, 50 µL of SmLcar** and 50 µL of a hundred-fold diluted sample of SmLcar_GY* cells). The main culture was incubated at 37 °C while shaking at 135 rpm until an $OD_{600}$ of 0.4 was reached. At this point, gene expression was induced by addition of IPTG (final concentration 0.1 mM) and cells were incubated at 37 °C while shaking at 135 rpm for 3-4 more hours, until $OD_{600}$ ~0.8. During this incubation step, selection medium was freshly prepared containing 25 µg/mL spectinomycin, 17.5 µg/mL chloramphenicol, 0.1 mM IPTG, 125 µM $MnCl_2$, 500 µM cam-3nY, and 20 µg/mL carbenicillin. The induced culture was then diluted 1:100 in 5 mL selection medium and grown for 24-48 hours at 30 °C and 135 rpm, while routinely measuring $OD_{600}$. Upon reaching $OD_{600}$ > ~0.7, a 1:100 dilution was performed in fresh selection medium, keeping the carbenicillin concentration constant at 20 µg/ml. Two such serial passages were performed in selection medium. Prior to selection and after each serial passage, overnight cultures of the mixed populations were inoculated in LB with 50 µg/mL spectinomycin and 35 µg/ml chloramphenicol. Plasmids were isolated and the change in the composition of the population throughout the mock selection was visualized via restriction digest. In addition, variants in the population were identified by Sanger sequencing with *DuetDOWN1*. Base calls taken from raw sequencing data were used to calculate the relative distribution of bases at positions of interest.

**Restriction digest:** Plasmids isolated from mock selections throughout the selection process were subjected to restriction enzyme double digestion in order to visualize the composition of the population. The 20 µL reactions contained 250 ng plasmid DNA, 0.4 µL *EcoRI*-HF (8 units), 0.4 µL *XhoI* (8 units), 2 µL CutSmart 10X buffer (final concentration 1X), and was topped up with MilliQ. The reactions were incubated at 37 °C for 1 hour, after which they were mixed with an appropriate amount of DNA loading dye, and the entire mixture was loaded on a freshly poured 1% agarose gel with Roti GelStain as a staining reagent. GeneRuler 1 kB DNA ladder (4 µL) was used as a marker. Agarose gel electrophoresis was performed in fresh TAE buffer for 80 minutes at 60 V, which is a relatively long running time and low voltage to ensure crisp bands and release of excess dye. Separated DNA was visualized under UV light and imaged.

**Residue selection mutagenesis:** Two residues that were previously identified to be important for the carbamoylase activity of SmLcar were used again, i.e. Leu217 and Phe329[2]. Next, following visual inspection of the crystal structure of wildtype SmLcar (PDB accession ID: 8APZ) and SmLcar mutant L217G/F329C (PDB accession ID: 8AQ0), neighboring residues in proximity to the catalytic cavity but not essential for catalysis, were selected for mutagenesis as well, i.e. Gln215 and Gly327.

**Generation of NNK libraries:** Overlap extension PCR (oePCR) with primers bearing degenerate NNK codons was employed to randomize the previously identified positions. Starting from pACYC_SmLcar, three PCR fragments with partially overlapping ends were generated using different *NNK_for* mutagenic primers in combination with their respective *_rev* primers (Table below). *DpnI* digestion was performed after the first PCR to remove any remaining template plasmid. Because coupling of more than two fragments using oePCR directly did not yield the desired full-length product, an intermediate fragment was generated out of two fragments using an additional PCR reaction, in the presence of the appropriate

*NNK_for* and *SmLcarCDS_GG_rev* primers (Table below). In the final step, this intermediate fragment was amplified together with its equivalent partially-overlapping fragment and the full-length, 1280 bp oePCR products were obtained by performing the reactions in presence of both *SmLcarCDS_GG_for* and *_rev* primers (Table below).

| **Lib_2N** | **Template** | **Primer pair** | **Fragment size** |
|---|---|---|---|
| Fragment 2posA | pACYC_SmLcar | *SmLcarCDS_GG_for + L217_rev* | 662 bp |
| Fragment 2posB | pACYC_SmLcar | *L217NNK_for + F329_rev* | 353 bp |
| Fragment 2posC | pACYC_SmLcar | *F329NNK_for + SmLcarCDS_GG_rev* | 299 bp |
| Intermediate fragment 2posBC | Fragment 2posB and 2posC, partially overlapping | *L217NNK_for + SmLcarCDS_GG_rev* | 634 bp |
| Full-length insert | Fragment 2posA and intermediate fragment 2posBC, partially overlapping | *SmlcarCDS_GG_for + SmLcarCDS_GG_rev* | 1280 bp |
| **Lib_4N** | **Template** | **Primer pair** | **Fragment size** |
| Fragment 4posA | pACYC_SmLcar | *SmLcarCDS_GG_for + Q215_L217_rev* | 656 bp |
| Fragment 4posB | pACYC_SmLcar | *Q215NNK_L217NNK_for + G327_F329_rev* | 355 bp |
| Fragment 4posC | pACYC_SmLcar | *G327NNK_F329NNK_for + SmLcarCDS_GG_rev* | 305 bp |
| Intermediate fragment 4posBC | Fragment 4posA and intermediate fragment 4posBC, partially overlapping | *Q215NNK_L217NNK_for + SmLcarCDS_GG_rev* | 642 bp |
| Full-length insert | Fragment 4posA and intermediate fragment 4posBC, partially overlapping | *SmlcarCDS_GG_for + SmLcarCDS_GG_rev* | 1280 bp |

In total, three PCR reactions were necessary for each library to create the full-length insert. The following PCR protocol was used: (1) initial denaturation at 95 ˚C for 3 min, (2) 30 cycles of denaturation at 95 ˚C for 30 s, annealing at 60˚C for 30 s and extension at 72 ˚C for 30 s; (3) a final extension at 72 ˚C for 10 min. After each PCR, the fragment of interest was excised from a 0.8% agarose gel and purified. All PCR reactions were performed in Milli-Q water with Phusion-HF DNA polymerase, 200 µM dNTPs, 0.25 µM forward and 0.25 µM reverse primer, 1X GC buffer, 3% DMSO, and ~1-5 ng template DNA in 50 µL reactions.

To assemble the selection plasmids, the partially-randomized CDS was cloned in pACYC_TEM-1.B9 using Golden Gate Assembly with Hi-T4 DNA ligase and *BsaI*-HF. The thermocycler program used for these assemblies was as follows: (1) 30 cycles alternating between 37 ˚C and 16 ˚C for 5 and 10 min, respectively, (2) a final digestion step at 55 ˚C for 10 min and (3) an enzyme inactivation step at 65 ˚C for 20 min.

The restriction-ligation reactions were initially transformed into NEB 10-beta cells. For Lib_2N, chemically competent NEB 10-beta cells were transformed (heat shock). For Lib_4N, freshly made electrocompetent NEB 10-beta cells were transformed[4] in order to increase transformation efficiency to accommodate the increased library size. The transformation efficiency was calculated based on the number of colony-forming units and final culture volume (**Supplementary Table S1**). All colonies were scraped from large selective LB agar plates (~50 mL) containing chloramphenicol (35 µg/ml). Plasmid DNA was isolated and sequenced with *DuetDOWN1* to verify library quality. Base calls taken from raw sequencing data were used to calculate the relative distribution of bases at the NNK positions. The isolated selection plasmids containing 2N or 4N libraries were stored at -20 ˚C until the start of a selection experiment.

**Selections of improved SmLcar variants from large libraries in liquid media:** Chemically competent or freshly prepared electrocompetent *E. coli* BL21(DE3) bearing pULTRA_3iY were transformed with selections plasmids containing either Lib_2N or Lib_4N. After recovery, all transformants were immediately grown as a preculture in 4 or 5 mL SOC medium containing 50 µg/mL spectinomycin and 35 µg/ml chloramphenicol, and a fraction of the cells was plated on LB agar plates with the same antibiotics to calculate the transformation efficiency for Lib_4N (**Supplementary Table S1**). After overnight shaking at 37 °C, main cultures of 5 mL LB containing 25 µg/mL spectinomycin and 17.5 µg/mL chloramphenicol were started with 50 µL (for Lib_2N) or 100 µL (for Lib_4N) of the densely grown overnight preculture. Multiple main cultures were started from the same preculture to create replicate populations and explore multiple evolutionary trajectories in parallel, i.e. Lib_2N was used to start two populations (2N_A, 2N_B) and Lib_4N was used to start four populations (4N_A, 4N_B, 4N_C, and 4N_D). The main cultures were incubated at 37 °C while shaking at 135 rpm until an $OD_{600}$ of 0.3 was reached. At this point, gene expression was induced by addition of IPTG (final concentration 0.1 mM) and cells were incubated at 37 °C while shaking at 135 rpm for 3 more hours, until $OD_{600}$ ~1. During this incubation step, selection medium was freshly prepared containing 25 μg/mL spectinomycin, 17.5 μg/mL chloramphenicol, 0.1 mM IPTG, 125 μM $MnCl_2$, 500 μM cam-3nY, and 20 μg/mL carbenicillin.

To start the selections, the induced cultures were diluted 1:100 in 5 mL selection medium, except for the first selection round of Lib_4N, which was diluted 1:25 to ensure sufficient representation of the library size. Selection cultures were grown at 30 °C and 135 rpm, while routinely measuring $OD_{600}$ using a portable spectrophotometer. Upon reaching sufficiently high cell density ($OD_{600}$ > ~0.6), a 1:100 dilution was performed in fresh selection medium. This was usually after 20-30 hours, with some cultures needing as little as 17 hours or as much as 42 hours depending on the selection progress and stringency. The selection stringency was occasionally increased by raising the concentration of carbenicillin from 20, to 50, to 100 μg/mL carbenicillin. As such, several growth-dilution passages were performed while monitoring the change in the composition of the population by sequencing (see below) as weaker variants are outcompeted by stronger ones. 3-4 selection rounds were performed for the 2N libraries and 5-6 rounds for the 4N libraries. It should be noted that 4N_B required >48 hours to reach a sufficiently high $OD_{600}$ and was discontinued, as the carbenicillin pressure would become less reliable after this time.

After each passage, a 25% glycerol stock of the mixed population was made to enable its long-term storage, and from these stocks the selection process could be restarted and continued. For 4N_A and 4N_D, selection cultures were restarted as such in order to increase the number of evolutionary trajectories and to diversify the possible outcomes.

The change in the composition of the populations was monitored by sequencing of the SmLcar gene. Prior to selection and after each serial passage, overnight cultures of the mixed populations were inoculated in LB with 50 μg/mL spectinomycin and 35 μg/ml chloramphenicol. Additionally, random candidates (single colonies) prior to selection and at the end of each selection campaign were inoculated in the same manner. Plasmids were isolated and variants in the population were identified by Sanger sequencing with *DuetDOWN1*. For mixed populations, base calls taken from raw sequencing data were used to calculate the relative distribution of bases at positions of interest. Once the composition of each population did not change anymore, they were considered to have sufficiently converged and the selections were finished up to identify the remaining variants.

**Chemical complementation screening in 96-well plates:** This protocol is based on the screening in liquid media as performed in previous work.[2] 96-deep well plates filled with LB medium (0.5 mL) containing 25 μg/mL spectinomycin and 17.5 μg/mL chloramphenicol were inoculated with single colonies from the agar plates of the starting or surviving library members from the selection campaigns, i.e. *E. coli* BL21(DE3) cells harboring pULTRA-3iY and a mutagenized SmLcar CDS cloned into pACYC_TEM-1.B9. In addition to these library members, a single colony of SmLcar_Leu217Gly-Phe329Tyr-Ala346Thr (SmLcar_GY), cloned into pACYC_TEM-1.B9, was used as a positive control. The resulting 96-deep well plates were incubated overnight at 37 °C while shaking at 900 rpm (*Titramax 1000 & Incubator 1000, Heidolph*). The next morning, 10 μL of the densely grown overnight cultures was transferred into fresh 96-deep well plates containing 0.5 mL LB media and 25 μg/mL spectinomycin and 17.5 μg/mL chloramphenicol. Additionally, some cells of each well were stamped onto LB agar plates with the same antibiotics and cultured at 37 °C overnight, for later isolation of plasmids. The bacteria in the 96-deep well plates were cultured at 37 °C for 1.5 hours while shaking at 900 rpm. Subsequently, expression was induced by addition of 16.5 µL of LB with appropriate antibiotics and 3 mM IPTG (final concentration of 0.1 mM IPTG) and plates were incubated at 37 °C for 3 more hours while shaking (900 rpm). Two transparent 96-well assay plates were set up in parallel by adding in each well: (1) 190 µL of LB with 25 µg/mL spectinomycin, 17.5 µg/mL chloramphenicol, 0.1 mM IPTG, 125 µM $MnCl_2$, 500 µM cam-3nY, and either 526 µg/µL or 105 µg/mL carbenicillin (final concentration 500 or 100 µg/mL, respectively) and (2) 10 µl of induced main culture ($OD_{600}$ ~1). Next, the two assay plates were transferred into two Synergy H1 microplate readers (*BioTek*) that were preheated to 30 °C. $OD_{600}$ was measured every 10 minutes from the bottom of the wells for 48 hours while continuously shaking (double orbital, 425 c.p.m.). The SmLcar gene variant featured in cells with the highest growth rate were identified by sequencing with *DuetDOWN1* after overnight growth in LB with appropriate antibiotics and plasmid isolation.

The CDS of the SmLcar gene variants of hits were cloned out by standard PCR with primers *SmLcarCDS_GG_for* and *SmLcarCDS_GG_rev*, and recloned into clean pACYC_TEM-1.B9 vectors by Golden Gate Assembly (as described earlier) to remove any possible spontaneous mutations in the backbone . Then, these variants were screened in 96-well plates as described above using a range of carbenicillin concentrations (50, 100, 150, 200 and 500 µg/ml), using SmLcar_Leu217Gly as a control for a slow-growing phenotype. Additionally, positive controls were included at 500 µg/ml carbenicillin which were supplied with 500 µM 3nY (instead of 500 µM cam-3nY) for each SmLcar variant.

**Categorization of library members as fast, medium, or slow growers:** Based on the $OD_{600}$ values over time as recorded by the abovementioned plate reader screenings, library members were categorized as fast, medium, or slow growers, aided by the phenotype of positive control SmLcar_GY as a fast-growing reference. Specifically, the growth rate is examined under high or moderate selective pressures (500 µg/ml or 100 µg/ml carbenicillin) and the growth rates of each library member is categorized as (1) fast, if the $OD_{600}$ has reached or has exceeded 0.3 after 24 hours; (2) slow, if the $OD_{600}$ has exceeded 0.3 after 48 hours; or (3) no growth, if the $OD_{600}$ failed to exceed 0.3 after 48 hours.

**Production and purification of SmLcar variants:** The production and purification of SmLcar variants is based on our previously described proceedings with minor adjustments.[2] The CDS of the SmLcar gene variants of hits was cloned by standard PCR with primers *SmLcarCDS_GG_for* and *SmLcarCDS_GG_rev*, and reinserted into clean pACYC_GG expression vectors by Golden Gate Assembly (as described earlier) to prepare them for purification. Flasks containing 250 mL or 500 mL LB medium with 35 µg/mL chloramphenicol were inoculated with 250 µL or 500 µL, respectively, of a densely grown overnight culture of *E. coli* BL21(DE3) cells harboring the appropriate pACYC_SmLcar plasmid. Cells were incubated at 37 °C and 135 rpm until an $OD_{600}$ of ~0.4 was reached and gene expression was induced by adding IPTG (final concentration 1 mM). Enzymes were produced for 4 hours at 37 °C while shaking (135 rpm), after which cells were harvested by centrifugation (3,700 rpm for 10 min, 4 °C). Cell pellets were resuspended in buffer (20 mL, 50 mM $Na_2HPO_4$, 500 mM NaCl, pH 8, containing 1 mg/mL egg white lysozyme. The cells were then lysed by sonication (10 min, 5 s pulse and pause cycles, 70% amplitude), and cellular debris was removed by centrifugation (12,000 rcf for 45 min, 4 °C). The supernatant was loaded onto a Ni-NTA resin and purified according to the manufacturer's specifications. Protein-containing fractions were pooled after elution, concentrated, and finally stored in 50 mM $Na_2HPO_4$, 150 mM NaCl, pH 7 with 5% glycerol at -20 °C. The purity and identity of SmLcar variants was confirmed by SDS-

PAGE (**Supplementary Fig. S1)** and mass spectrometry, respectively (Q-TOF MS, **Supplementary Table S8**).

**In vitro characterization of carbamoylases:** This protocol was adapted from previous work.[2] Standard enzymatic reactions were performed in 50 mM $Na_2HPO_4$, 150 mM NaCl buffer with purified SmLcar variants in 400 µL reaction volume, supplemented with 1 $MnCl_2$. The pH of the buffer was different depending on the substrate: either 8.0 (with 2 mM of cam-3iY or cam-Y), or 6.0 (with 2 mM cam-3nY). Substrates were prepared as 10x stocks. To characterize the evolved variants, 40x stock solutions of the enzyme were prepared in the same buffer. To initiate the reactions, 10 µL of the 40 µM stock solution of each evolved variant was added to the mixture (giving a final enzyme concentration of 1 µM). For the wildtype, a final enzyme concentration of 10 µM was used instead. The reactions were performed in triplicate and incubated at room temperature (25 °C) without shaking, and aliquots thereof were quenched by addition of one volume equivalent of 3% $H_3PO_4$. After centrifugation (10 min, 15060 rcf), the supernatants were analyzed by reverse-phase HPLC without further dilution or purification. The concentrations of cam-(nc)AAs and (nc)AAs were determined based on the area under the curve of the substrate and product peak, as described in the Materials & Methods section (see the calibration curves in **Supplementary Fig. S2**). The initial rate of each enzyme was calculated based on the appearance of (nc)AA after 5 minutes, 15 minutes, or 30 minutes, depending on each enzyme's activity. For the wildtype, which has a much lower activity, a 5-hour or 24-hour timepoint was used.

#### 4. Sequences

Sequences of primers used in this work, with mutations or randomizations shown underlined and bold.

| Name | Sequence (5' → 3') |
|---|---|
| DuetUP2 | TTGTACACGGCCGCATAATC |
| DuetDOWN1 | GATTATGCGGCCGTGTACAA |
| MCS1_Up | GGAGATATACCATGGGCAGC |
| pACYC_GG_3rev | GGGCAATCAGCTGTTGCCCGTTTCACTGGTGAAAAGAAAAAC |
| SmLcarCDS_GG_for | ACCGGTCTCCTGGTATGGCGGCGCCTGGT |
| SmLcarCDS_GG_rev | GTTGGTCTCCCAAGTCACTCCACAATCTC |
| L217NNK_for | TCACTCATTGTCAAGGC**NNK**TGGTGGCTGGAATTTACTTTGAC |
| Q215NNK_L217NNK_for | TGGCGTTGTCACTCATTGT**NNK**GGC**NNK**TGGTGGCTGGAATTTAC |
| L217_rev | GCCTTGACAATGAGTGACAACGCCAATTTG |
| Q215_L217_rev | ACAATGAGTGACAACGCCAATTTGCTTATTC |
| G327NNK_F329NNK_for | TGCTCGATTGAAGCAGTA**NNK**CAC**NNK**GACCCAGTAACCTTTG |
| G327_F329_rev | TACTGCTTCAATCGAGCATCCGACACCCAGAC |
| SmLcar_TTT-TTC_F222F_fw | GGTGGCTGGAATT**C**ACTTTGACAGGTCGCGAGG |
| SmLcar_TTT-TTC_F222F_rv | CCTCGCGACCTGTCAAAGT**G**AATTCCAGCCACC |
| SmLcar_GTC-CTC_QC_fw | GGCTCTCAAGGGCATCG**C**TCGAGATCCCGGTGCCT |
| SmLcar_GTC-CTC_QC_rv | CGATGCCCTTGAGAGCCTTCAACC |
| SmLcar_M141I_fw | GCTTTGCACCTGCAAT**A**TTGGCCTCTGGCGTATT |
| SmLcar_M141_rev | ATTGCAGGTGCAAAGCGCGCCCCCT |

**Sequence of wildtype SmLcar CDS (5' → 3')**

An N-terminal His-tag, spacer, and a *BsaI* scar precede the SmLcar CDS and are underlined.

<u>ATGGGCAGCAGCCATCATCATCATCATCACGGCAGCGGCCTGGTGCCGCGCGGCAGCGCTGGT</u>ATGGCGGCGCCT
GGTGAAAACCGTCGTGTAAATGCCGATCGCTTATGGGACTCGCTGATGGAAATGGCGAAAATTGGCCCTGGAGTT
GCGGGAGGTAACAACCGCCAGACCTTGACAGACGCTGACGGGGAAGGGCGCCGCCTGTTTCAATCCTGGTGTGAA
GAAGCCGGCTTGTCGATGGGCGTCGATAAGATGGGCACCATGTTTCTTACACGCCCTGGAACTGACCCCGACGCA
TTGCCAGTTCATATTGGCTCGCACCTGGATACTCAGCCTACCGGAGGAAAATTTGACGGCGTCTTAGGTGTATTG
AGCGGACTGGAAGCTGTCCGTACGATGAATGACTTGGGAATCAAGACCAAACACCCAATTGTCGTTACCAATTGG
ACTAACGAGGAGGGGGCGCGCTTTGCACCTGCAATGTTGGCCTCTGGCGTATTTGCAGGCGTACACACCTTAGAA
TATGCCTACGCCCGTAAGGACCCAGAAGGCAAATCTTTCGGTGATGAATTAAAGCGTATCGGCTGGTTGGGCGAT
GAGGAAGTTGGTGCGCGTAAAATGCACGCATACTTTGAATACCACATCGAGCAAGGCCCAATTCTTGAGGCGGAG
AATAAGCAAATTGGCGTTGTCACTCATTGTCAAGGCCTGTGGTGGCTGGAATTTACTTTGACAGGTCGCGAGGCT
CATACCGGATCAACACCGATGGATATGCGTGTGAATGCGGGATTAGCGATGGCTCGTATTCTTGAAATGGTCCAA
ACCGTTGCCATGGAGAACCAACCCGGTGCAGTGGGCGGCGTGGGGCAGATGTTTTTTAGCCCTAACTCCCGCAAC
GTGTTACCGGGAAAAGTAGTTTTCACAGTAGACATCCGTTCCCCTGATCAAGCAAAGCTGGATGGAATGCGCGCA
CGCATTGAGGCAGAAGCGCCAAAAATTTGTGAGCGTCTGGGTGTCGGATGCTCGATTGAAGCAGTAGGTCACTTC
GACCCAGTAACCTTTGACCCTAAACTTGTAGAGACAGTTCGTGGGGCGGCAGAGAAATTAGGGTATTCGCATATG
AACTTAGTTTCTGGCGCCGGACACGACGCCTGCTGGGCGGCTAAGGTTGCACCTACGACAATGATCATGTGTCCT
TGTGTGGGCGGACTTTCGCATAATGAGGCGGAAGATATTTCTCGTGAATGGGCAGCAGCCGGTGCAGACGTCCTT
TTTCATGCTGTTTTGGAAACCGCTGAGATTGTGGAGTGA

#### Sequence of TEM-1.B9 CDS (5’ → 3’)

Suppressed stop codon (where the ncAA is incorporated) shown in bold red. A spacer precedes the TEM-1.B9 CDS and is underlined.

ATGTTCGGCAGCATGAGTATTCAACATTTCCGTGTCGCCCTTATTCCCTTTTTTGCGGCATTTTGCCTTCCTGTT
TTTGCTCACCCAGAAACGCTGGTGAAAGTAAAAGATGCTGAAGATCAGTTGGGTGCACGAGTGGGTTACATCGAA
CTGGATCTCAACAGCGGTAAGATCCTTGAGAGTTTTCGCCCCGAAGAACGTTTTCCATTCATGAGCACTGGGAAA
GTTCTGCTATGTGGCGCGGTATTATCCCGTGTTGACGCCGGGCAAGAGCAACTCGGTCGCCGCATACACTATTCT
CAGAATGACTTGGTTGAGTACTCACCAGTCACAGAAAAGCATCTTACGGATGGCATGACAGTAAGAGAATTATGC
AGTGCTGCCATAACCATGAGTGATAACACTGCGGCCAACTTACTTACCACAACGATCGGAGGACCGAAGGAGCTC
ACCGCTTTTATGCACAACATGGGGGATCATGTAACTCGC**TAG**GATCGTTGGGAACCGGAGTTGAATGAAGCCATA
CCAAACGACGAGCGTGACACCACGATGCCTGCAGCAATGGCAACAACGTTGCGCAAACTATTAACTGGCGAACTA
CTTACTCTAGCTTCCCGGCAACAATTAATAGACTGGATGGAGGCGGATAAAGTTGCAGGACCACTTCTGCGCTCG
GCCCTTCCGGCTGGCTGGTTTATTGCTGATAAATCTGGAGCCGGTGAGCGTGGGTCcCGCGGTATCATTGCAGCA
CTGGGGCCAGATGGTAAGCCCTCCCGTATCGTAGTTATCTACACGACGGGGAGTCAGGCAACTATGGATGAACGA
AATAGACAGATCGCTGAGATAGGTGCCTCACTGATTAAGCATTGGTAA

#### Sequence of all the SmLcar hits

His-tags, spacers, and *BsaI* scars are left out, and only the SmLcar CDS of each respective hit is given. Deviations from the wildtype sequence and NNK positions are shown underlined and bold. Some sequences have additional mutations outside of the designed NNK positions, as a consequence of the random errors introduced by PCR or cell division.

#### SmLcar_GY or SmLcar_L217G-F329Y-A346T

ATGGCGGCGCCTGGTGAAAACCGTCGTGTAAATGCCGATCGCTTATGGGACTCGCTGATGGAAATGGCGAAAATT
GGCCCTGGAGTTGCGGGAGGTAACAACCGCCAGACCTTGACAGACGCTGACGGGGAAGGGCGCCGCCTGTTTCAA
TCCTGGTGTGAAGAAGCCGGCTTGTCGATGGGCGTCGATAAGATGGGCACCATGTTTCTTACACGCCCTGGAACT
GACCCCGACGCATTGCCAGTTCATATTGGCTCGCACCTGGATACTCAGCCTACCGGAGGAAAATTTGACGGCGTC
TTAGGTGTATTGAGCGGACTGGAAGCTGTCCGTACGATGAATGACTTGGGAATCAAGACCAAACACCCAATTGTC
GTTACCAATTGGACTAACGAGGAGGGGGCGCGCTTTGCACCTGCAATGTTGGCCTCTGGCGTATTTGCAGGCGTA
CACACCTTAGAATATGCCTACGCCCGTAAGGACCCAGAAGGCAAATCTTTCGGTGATGAATTAAAGCGTATCGGC
TGGTTGGGCGATGAGGAAGTTGGTGCGCGTAAAATGCACGCATACTTTGAATACCACATCGAGCAAGGCCCAATT
CTTGAGGCGGAGAATAAGCAAATTGGCGTTGTCACTCATTGTCAAGGC**GGG**TGGTGGCTGGAATTTACTTTGACA
GGTCGCGAGGCTCATACCGGATCAACACCGATGGATATGCGTGTGAATGCGGGATTAGCGATGGCTCGTATTCTT
GAAATGGTCCAAACCGTTGCCATGGAGAACCAACCCGGTGCAGTGGGCGGCGTGGGGCAGATGTTTTTTAGCCCT
AACTCCCGCAACGTGTTACCGGGAAAAGTAGTTTTCACAGTAGACATCCGTTCCCCTGATCAAGCAAAGCTGGAT
GGAATGCGCGCACGCATTGAGGCAGAAGCGCCAAAAATTTGTGAGCGTCTGGGTGTCGGATGCTCGATTGAAGCA
GTAGGTCAC**TAT**GACCCAGTAACCTTTGACCCTAAACTTGTAGAGACAGTTCGTGGGGCG**A**CAGAGAAATTAGGG
TATTCGCATATGAACTTAGTTTCTGGCGCCGGACACGACGCCTGCTGGGCGGCTAAGGTTGCACCTACGACAATG
ATCATGTGTCCTTGTGTGGGCGGACTTTCGCATAATGAGGCGGAAGATATTTCTCGTGAATGGGCAGCAGCCGGT
GCAGACGTCCTTTTTCATGCTGTTTTGGAAACCGCTGAGATTGTGGAGTGA

#### SmLcar_GL or SmLcar_M141I-L217G-F329L

ATGGCGGCGCCTGGTGAAAACCGTCGTGTAAATGCCGATCGCTTATGGGACTCGCTGATGGAAATGGCGAAAATT
GGCCCTGGAGTTGCGGGAGGTAACAACCGCCAGACCTTGACAGACGCTGACGGGGAAGGGCGCCGCCTGTTTCAA
TCCTGGTGTGAAGAAGCCGGCTTGTCGATGGGCGTCGATAAGATGGGCACCATGTTTCTTACACGCCCTGGAACT
GACCCCGACGCATTGCCAGTTCATATTGGCTCGCACCTGGATACTCAGCCTACCGGAGGAAAATTTGACGGCGTC
TTAGGTGTATTGAGCGGACTGGAAGCTGTCCGTACGATGAATGACTTGGGAATCAAGACCAAACACCCAATTGTC
GTTACCAATTGGACTAACGAGGAGGGGGCGCGCTTTGCACCTGCAAT**A**TTGGCCTCTGGCGTATTTGCAGGCGTA
CACACCTTAGAATATGCCTACGCCCGTAAGGACCCAGAAGGCAAATCTTTCGGTGATGAATTAAAGCGTATCGGC
TGGTTGGGCGATGAGGAAGTTGGTGCGCGTAAAATGCACGCATACTTTGAATACCACATCGAGCAAGGCCCAATT
CTTGAGGCGGAGAATAAGCAAATTGGCGTTGTCACTCATTGTCAAGGC**GGG**TGGTGGCTGGAATTTACTTTGACA
GGTCGCGAGGCTCATACCGGATCAACACCGATGGATATGCGTGTGAATGCGGGATTAGCGATGGCTCGTATTCTT
GAAATGGTCCAAACCGTTGCCATGGAGAACCAACCCGGTGCAGTGGGCGGCGTGGGGCAGATGTTTTTTAGCCCT
AACTCCCGCAACGTGTTACCGGGAAAAGTAGTTTTCACAGTAGACATCCGTTCCCCTGATCAAGCAAAGCTGGAT
GGAATGCGCGCACGCATTGAGGCAGAAGCGCCAAAAATTTGTGAGCGTCTGGGTGTCGGATGCTCGATTGAAGCA
GTAGGTCAC**CTT**GACCCAGTAACCTTTGACCCTAAACTTGTAGAGACAGTTCGTGGGGCGGCAGAGAAATTAGGG
TATTCGCATATGAACTTAGTTTCTGGCGCCGGACACGACGCCTGCTGGGCGGCTAAGGTTGCACCTACGACAATG
ATCATGTGTCCTTGTGTGGGCGGACTTTCGCATAATGAGGCGGAAGATATTTCTCGTGAATGGGCAGCAGCCGGT
GCAGACGTCCTTTTTCATGCTGTTTTGGAAACCGCTGAGATTGTGGAGTGA

#### SmLcar_DI or SmLcar_M141I-L217D-Q296R-F329I

ATGGCGGCGCCTGGTGAAAACCGTCGTGTAAATGCCGATCGCTTATGGGACTCGCTGATGGAAATGGCGAAAATT
GGCCCTGGAGTTGCGGGAGGTAACAACCGCCAGACCTTGACAGACGCTGACGGGGAAGGGCGCCGCCTGTTTCAA
TCCTGGTGTGAAGAAGCCGGCTTGTCGATGGGCGTCGATAAGATGGGCACCATGTTTCTTACACGCCCTGGAACT
GACCCCGACGCATTGCCAGTTCATATTGGCTCGCACCTGGATACTCAGCCTACCGGAGGAAAATTTGACGGCGTC
TTAGGTGTATTGAGCGGACTGGAAGCTGTCCGTACGATGAATGACTTGGGAATCAAGACCAAACACCCAATTGTC
GTTACCAATTGGACTAACGAGGAGGGGGCGCGCTTTGCACCTGCAAT**A**TTGGCCTCTGGCGTATTTGCAGGCGTA
CACACCTTAGAATATGCCTACGCCCGTAAGGACCCAGAAGGCAAATCTTTCGGTGATGAATTAAAGCGTATCGGC
TGGTTGGGCGATGAGGAAGTTGGTGCGCGTAAAATGCACGCATACTTTGAATACCACATCGAGCAAGGCCCAATT
CTTGAGGCGGAGAATAAGCAAATTGGCGTTGTCACTCATTGTCAAGGC**GAT**TGGTGGCTGGAATTTACTTTGACA
GGTCGCGAGGCTCATACCGGATCAACACCGATGGATATGCGTGTGAATGCGGGATTAGCGATGGCTCGTATTCTT
GAAATGGTCCAAACCGTTGCCATGGAGAACCAACCCGGTGCAGTGGGCGGCGTGGGGCAGATGTTTTTTAGCCCT
AACTCCCGCAACGTGTTACCGGGAAAAGTAGTTTTCACAGTAGACATCCGTTCCCCTGATC**G**AGCAAAGCTGGAT
GGAATGCGCGCACGCATTGAGGCAGAAGCGCCAAAAATTTGTGAGCGTCTGGGTGTCGGATGCTCGATTGAAGCA
GTAGGTCAC**ATT**GACCCAGTAACCTTTGACCCTAAACTTGTAGAGACAGTTCGTGGGGCGGCAGAGAAATTAGGG
TATTCGCATATGAACTTAGTTTCTGGCGCCGGACACGACGCCTGCTGGGCGGCTAAGGTTGCACCTACGACAATG
ATCATGTGTCCTTGTGTGGGCGGACTTTCGCATAATGAGGCGGAAGATATTTCTCGTGAATGGGCAGCAGCCGGT
GC**G**GACGTCCTTTTTCATGCTGTTTTGGAAACCGCTGAGATTGTGGAGTGA

#### SmLcar_IGTA or SmLcar_Q215I-L217G-G327T-F329A-A368T

ATGGCGGCGCCTGGTGAAAACCGTCGTGTAAATGCCGATCGCTTATGGGACTCGCTGATGGAAATGGCGAAAATT
GGCCCTGGAGTTGCGGGAGGTAACAACCGCCAGACCTTGACAGACGCTGACGGGGAAGGGCGCCGCCTGTTTCAA
TCCTGGTGTGAAGAAGCCGGCTTGTCGATGGGCGTCGATAAGATGGGCACCATGTTTCTTACACGCCCTGGAACT
GACCCCGACGCATTGCCAGTTCATATTGGCTCGCACCTGGATACTCAGCCTACCGGAGGAAAATTTGACGGCGTC
TTAGGTGTATTGAGCGGACTGGAAGCTGTCCGTACGATGAATGACTTGGGAATCAAGACCAAACACCCAATTGTC
GTTACCAATTGGACTAACGAGGAGGGGGCGCGCTTTGCACCTGCAATGTTGGCCTCTGGCGTATTTGCAGGCGTA
CACACCTTAGAATATGCCTACGCCCGTAAGGACCCAGAAGGCAAATCTTTCGGTGATGAATTAAAGCGTATCGGC
TGGTTGGGCGATGAGGAAGTTGGTGCGCGTAAAATGCACGCATACTTTGAATACCACATCGAGCAAGGCCCAATT
CTTGAGGCGGAGAATAAGCAAATTGGCGTTGTCACTCATTGT**ATT**GGC**GGG**TGGTGGCTGGAATTTACTTTGACA
GGTCGCGAGGCTCATACCGGATCAACACCGATGGATATGCGTGTGAATGCGGGATTAGCGATGGCTCGTATTCTT
GAAATGGTCCAAACCGTTGCCATGGAGAACCAACCCGGTGCAGTGGGCGGCGTGGGGCAGATGTTTTTTAGCCCT
AACTC**T**CGCAACGTGTTACCGGGAAAAGTAGTTTTCACAGTAGACATCCGTTCCCCTGATCAAGCAAAGCTGGAT
GGAATGCGCGCACGCATTGAGGCAGAAGCGCCAAAAATTTGTGAGCGTCTGGGTGTCGGATGCTCGATTGAAGCA
GTA**ACT**CAC**GCG**GACCCAGTAACCTTTGACCCTAAACTTGTAGAGACAGTTCGTGGGGCGGCAGAGAAATTAGGG
TATTCGCATATGAACTTAGTTTCTGGCGCCGGACACGACGCCTGCTGGGCG**A**CTAAGGTTGCACCTACGACAATG
ATCATGTGTCCTTGTGTGGGCGGACTTTCGCATAATGAGGCGGAAGATATTTCTCGTGAATGGGCAGCAGCCGGT
GCAGACGTCCTTTTTCATGCTGTTTTGGAAACCGCTGAGATTGTGGAGTGA

**SmLcar_TGTA or SmLcar_Q215T-L217G-G327T-F329A**

ATGGCGGCGCCTGGTGAAAACCGTCGTGTAAATGCCGATCGCTTATGGGACTCGCTGATGGAAATGGCGAAAATT
GGCCCTGGAGTTGCGGGAGGTAACAACCGCCAGACCTTGACAGACGCTGACGGGGAAGGGCGCCGCCTGTTTCAA
TCCTGGTGTGAAGAAGCCGGCTTGTCGATGGGCGTCGATAAGATGGGCACCATGTTTCTTACACGCCCTGGAACT
GACCCCGACGCATTGCCAGTTCATATTGGCTC**A**CACCTGGATACTCAGCCTACCGGAGGAAAATTTGACGGCGTC
TTAGGTGTATTGAGCGGACTGGAAGCTGTCCGTACGATGAATGACTTGGGAATCAAGACCAAACACCCAATTGTC
GTTACCAATTGGACTAACGAGGAGGGGGCGCGCTTTGCACCTGCAATGTTGGCCTCTGGCGTATTTGCAGGCGTA
CACACCTTAGAATATGCCTACGCCCGTAAGGACCCAGAAGGCAAATCTTTCGGTGATGAATTAAAGCGTATCGGC
TGGTTGGGCGATGAGGAAGTTGGTGCGCGTAAAATGCACGCATACTTTGAATACCACATCGAGCAAGGCCCAATT
CTTGAGGCGGAGAATAAGCAAATTGGCGTTGTCACTCATTGT**ACG**GGC**GGG**TGGTGGCTGGAATTTACTTTGACA
GGTCGCGAGGCTCATACCGGATCAACACCGATGGATATGCGTGTGAATGCGGGATTAGCGATGGCTCGTATTCTT
GAAATGGTCCAAACCGTTGCCATGGAGAACCAACCCGGTGCAGTGGGCGGCGTGGGGCAGATGTTTTTTAGCCCT
AACTCCCGCAACGTGTTACCGGGAAAAGTAGTTTTCACAGTAGACATCCGTTCCCCTGATCAAGCAAAGCTGGAT
GGAATGCGCGCACGCATTGAGGCAGAAGCGCCAAAAATTTGTGAGCGTCTGGGTGTCGGATGCTCGATTGAAGCA
GTA**ACG**CAC**GCG**GACCCAGTAACCTTTGACCCTAAACTTGTAGAGACAGTTCGTGGGGCGGCAGAGAAATTAGGG
TATTCGCATATGAACTTAGTTTCTGGCGCCGGACACGACGCCTGCTGGGCGGCTAAGGTTGCACCTACGACAATG
ATCATGTGTCCTTGTGTGGGCGGACTTTCGCATAATGAGGCGGAAGATATTTCTCGTGAATGGGCAGCAGCCGGT
GCAGACGTCCTTTTTCATGCTGTTTTGGAAACCGCTGAGATTGTGGAGTGA

**SmLcar_WTTS or SmLcar_ Q215W-L217T-G327T-F329S**

ATGGCGGCGCCTGGTGAAAACCGTCGTGTAAATGCCGATCGCTTATGGGACTCGCTGATGGAAATGGCGAAAATT
GGCCCTGGAGTTGCGGGAGGTAACAACCGCCAGACCTTGACAGACGCTGACGGGGAAGGGCGCCGCCTGTTTCAA
TCCTGGTGTGAAGAAGCCGGCTTGTCGATGGGCGTCGATAAGATGGGCACCATGTTTCTTACACGCCCTGGAACT
GACCCCGACGCATTGCCAGTTCATATTGGCTCGCACCTGGATACTCAGCCTACCGGAGGAAAATTTGACGGCGTC
TTAGGTGTATTGAGCGGACTGGAAGCTGTCCGTACGATGAATGACTTGGGAATCAAGACCAAACACCCAATTGTC
GTTACCAATTGGACTAACGAGGAGGGGGCGCGCTTTGCACCTGCAATGTTGGCCTCTGGCGTATTTGCAGGCGTA
CACACCTTAGAATATGCCTACGCCCGTAAGGACCCAGAAGGCAAATCTTTCGGTGATGAATTAAAGCGTATCGGC
TGGTTGGGCGATGAGGAAGTTGGTGCGCGTAAAATGCACGCATACTTTGAATACCACATCGAGCAAGGCCCAATT
CTTGAGGCGGAGAATAAGCAAATTGGCGTTGTCACTCATTGT**TGG**GGC**ACT**TGGTGGCTGGAATTTACTTTGACA
GGTCGCGAGGCTCATACCGGATCAACACCGATGGATATGCGTGTGAATGCGGGATTAGCGATGGCTCGTATTCTT
GAAATGGTCCAAACCGTTGCCATGGAGAACCAACCCGGTGCAGTGGGCGGCGTGGGGCAGATGTTTTTTAGCCCT
AACTCCCGCAACGTGTTACCGGGAAAAGTAGTTTTCACAGTAGACATCCGTTCCCCTGATCAAGCAAAGCTGGAT
GGAATGCGCGCACGCATTGAGGCAGAAGCGCCAAAAATTTGTGAGCGTCTGGGTGTCGGATGCTCGATTGAAGCA
GTA**ACG**CAC**AGT**GACCCAGTAACCTTTGACCCTAAACTTGTAGAGACAGTTCGTGGGGCGGCAGAGAAATTAGGG
TATTCGCATATGAACTTAGTTTCTGGCGCCGGACACGACGCCTGCTGGGCGGCTAAGGTTGCACCTACGACAATG
ATCATGTGTCCTTGTGTGGGCGGACTTTCGCATAATGAGGCGGAAGATATTTCTCGTGAATGGGCAGCAGCCGGT
GCAGACGTCCTTTTTCATGCTGTTTTGGAAACCGCTGAGATTGTGGAGTGA

**SmLcar_GGHR or SmLcar_ Q215G-L217G-G327H-F329R**

ATGGCGGCGCCTGGTGAAAACCGTCGTGTAAATGCCGATCGCTTATGGGACTCGCTGATGGAAATGGCGAAAATT
GGCCCTGGAGTTGCGGGAGGTAACAACCGCCAGACCTTGACAGACGCTGACGGGGAAGGGCGCCGCCTGTTTCAA
TCCTGGTGTGAAGAAGCCGGCTTGTCGATGGGCGTCGATAAGATGGGCACCATGTTTCTTACACGCCCTGGAACT
GACCCCGACGCATTGCCAGTTCATATTGGCTCGCACCTGGATACTCAGCCTACCGGAGGAAAATTTGACGGCGTC
TTAGGTGTATTGAGCGGACTGGAAGCTGTCCGTACGATGAATGACTTGGGAATCAAGACCAAACACCCAATTGTC
GTTACCAATTGGACTAACGAGGAGGGGGCGCGCTTTGCACCTGCAATGTTGGCCTCTGGCGTATTTGCAGGCGTA
CACACCTTAGAATATGCCTACGCCCGTAAGGACCCAGAAGGCAAATCTTTCGGTGATGAATTAAAGCGTATCGGC
TGGTTGGGCGATGAGGAAGTTGGTGCGCGTAAAATGCACGCATACTTTGAATACCACATCGAGCAAGGCCCAATT
CTTGAGGCGGAGAATAAGCAAATTGGCGTTGTCACTCATTGT**GGG**GGC**GGG**TGGTGGCTGGAATTTACTTTGACA
GGTCGCGAGGCTCATACCGGATCAACACCGATGGATATGCGTGTGAATGCGGGATTAGCGATGGCTCGTATTCTT
GAAATGGTCCAAACCGTTGCCATGGAGAACCAACCCGGTGCAGTGGGCGGCGTGGGGCAGATGTTTTTTAGCCCT
AACTCCCGCAACGTGTTACCGGGAAAAGTAGTTTTCACAGTAGACATCCGTTCCCCTGATCAAGCAAAGCTGGAT
GGAATGCGCGCACGCATTGAGGCAGAAGCGCCAAAAATTTGTGAGCGTCTGGGTGTCGGATGCTCGATTGAAGCA
GTA**CAT**CAC**CGG**GACCCAGTAACCTTTGACCCTAAACTTGTAGAGACAGTTCGTGGGGCGGCAGAGAAATTAGGG
TATTCGCATATGAACTTAGTTTCTGGCGCCGGACACGACGCCTGCTGGGCGGCTAAGGTTGCACCTACGACAATG
ATCATGTGTCCTTGTGTGGGCGGACTTTCGCATAATGAGGCGGAAGATATTTCTCGTGAATGGGCAGCAGCCGGT
GCAGACGTCCTTTTTCATGCTGTTTTGGAAACCGCTGAGATTGTGGAGTGA

#### SmLcar_AGHA or SmLcar_ Q215A-L217G-G327H-F329A-A368T

ATGGCGGCGCCTGGTGAAAACCGTCGTGTAAATGCCGATCGCTTATGGGACTCGCTGATGGAAATGGCGAAAATT
GGCCCTGGAGTTGCGGGAGGTAACAACCGCCAGACCTTGACAGACGCTGACGGGGAAGGGCGCCGCCTGTTTCAA
TCCTGGTGTGAAGAAGCCGGCTTGTCGATGGGCGTCGATAAGATGGGCACCATGTTTCTTACACGCCCTGGAACT
GACCCCGACGCATTGCCAGTTCATATTGGCTCGCACCTGGATACTCAGCCTACCGGAGGAAAATTTGACGGCGTC
TTAGGTGTATTGAGCGGACTGGAAGCTGTCCGTACGATGAATGACTTGGGAATCAAGACCAAACACCCAATTGTC
GTTACCAATTGGACTAACGAGGAGGGGGCGCGCTTTGCACCTGCAATGTTGGCCTCTGGCGTATTTGCAGGCGTA
CACACCTTAGAATATGCCTACGCCCGTAAGGACCCAGAAGGCAAATCTTTCGGTGATGAATTAAAGCGTATCGGC
TGGTTGGGCGATGAGGAAGTTGGTGCGCGTAAAATGCACGCATACTTTGAATACCACATCGAGCAAGGCCCAATT
CTTGAGGCGGAGAATAAGCAAATTGGCGTTGTCACTCATTGT**GCG**GGC**GGG**TGGTGGCTGGAATTTACTTTGACA
GGTCGCGAGGCTCATACCGGATCAACACCGATGGATATGCGTGTGAATGCGGGATTAGCGATGGCTCGTATTCTT
GAAATGGTCCAAACCGTTGCCATGGAGAACCAACCCGGTGCAGTGGGCGGCGTGGGGCAGATGTTTTTTAGCCCT
AACTCCCGCAACGTGTTACCGGGAAAAGTAGTTTTCACAGTAGACATCCGTTCCCCTGATCAAGCAAAGCTGGAT
GGAATGCGCGCACGCATTGAGGCAGAAGCGCCAAAAATTTGTGAGCGTCTGGGTGTCGGATGCTCGATTGAAGCA
GTA**CAT**CAC**GCT**GACCCAGTAACCTTTGACCCTAAACTTGTAGAGACAGTTCGTGGGGCGGCAGAGAAATTAGGG
TATTCGCATATGAACTTAGTTTCTGGCGCCGGACACGACGCCTGCTGGGCG**A**CTAAGGTTGCACCTACGACAATG
ATCATGTGTCCTTGTGTGGGCGGACTTTCGCATAATGAGGCGGAAGATATTTCTCGTGAATGGGCAGCAGCCGGT
GCAGACGTCCTTTTTCATGCTGTTTTGGAAACCGCTGAGATTGTGGAGTGA

#### SmLcar_AACA or SmLcar_ Q215A-L217A-G327C-F329A

ATGGCGGCGCCTGGTGAAAACCGTCGTGTAAATGCCGATCGCTTATGGGACTCGCTGATGGAAATGGCGAAAATT
GGCCCTGGAGTTGCGGGAGGTAACAACCGCCAGACCTTGACAGACGCTGACGGGGAAGGGCGCCGCCTGTTTCAA
TCCTGGTGTGAAGAAGCCGGCTTGTCGATGGGCGTCGATAAGATGGGCACCATGTTTCTTACACGCCCTGGAACT
GACCCCGACGCATTGCCAGTTCATATTGGCTCGCACCTGGATACTCAGCCTACCGGAGGAAAATTTGACGGCGTC
TTAGGTGTATTGAGCGGACTGGAAGCTGTCCGTACGATGAATGACTTGGGAATCAAGACCAAACACCCAATTGTC
GTTACCAATTGGACTAACGAGGAGGGGGCGCGCTTTGCACCTGCAATGTTGGCCTCTGGCGTATTTGCAGGCGTA
CACACCTTAGAATATGCCTACGCCCGTAAGGACCCAGAAGGCAAATCTTTCGGTGATGAATTAAAGCGTATCGGC
TGGTTGGGCGATGAGGAAGTTGGTGCGCGTAAAATGCACGCATACTTTGAATACCACATCGAGCAAGGCCCAATT
CTTGAGGCGGAGAATAAGCAAATTGGCGTTGTCACTCATTGT**GCG**GGC**GCT**TGGTGGCTGGAATTTACTTTGACA
GGTCGCGAGGCTCATACCGGATCAACACCGATGGATATGCGTGTGAATGCGGGATTAGCGATGGCTCGTATTCTT
GAAATGGTCCAAACCGTTGCCATGGAGAACCAACCCGGTGCAGTGGGCGGCGTGGGGCAGATGTTTTTTAGCCCT
AACTCCCGCAACGTGTTACCGGGAAAAGTAGTTTTCACAGTAGACATCCGTTCCCCTGATCAAGCAAAGCTGGAT
GGAATGCGCGCACGCATTGAGGCAGAAGCGCCAAAAATTTGTGAGCGTCTGGGTGTCGGATGCTCGATTGAAGCA
GTA**TGT**CAC**GCG**GACCCAGTAACCTTTGACCCTAAACTTGTAGAGACAGTTCGTGGGGCGGCAGAGAAATTAGGG
TATTCGCATATGAACTTAGTTTCTGGCGCCGGACACGACGCCTGCTGGGCGGCTAAGGTTGCACCTACGACAATG
ATCATGTGTCCTTGTGTGGGCGGACTTTCGCATAATGAGGCGGAAGATATTTCTCGTGAATGGGCAGCAGCCGGT
GCAGACGTCCTTTTTCATGCTGTTTTGGAAACCGCTGAGATTGTGGAGTGA

#### SmLcar_FGCA or SmLcar_ M141I-Q215F-L217G-G327C-F329A

ATGGCGGCGCCTGGTGAAAACCGTCGTGTAAATGCCGATCGCTTATGGGACTCGCTGATGGAAATGGCGAAAATT
GGCCCTGGAGTTGCGGGAGGTAACAACCGCCAGACCTTGACAGACGCTGACGGGGAAGGGCGCCGCCTGTTTCAA
TCCTGGTGTGAAGAAGCCGGCTTGTCGATGGGCGTCGATAAGATGGGCACCATGTTTCTTACACGCCCTGGAACT
GACCCCGACGCATTGCCAGTTCATATTGGCTCGCACCTGGATACTCAGCCTACCGGAGGAAAATTTGACGGCGTC
TTAGGTGTATTGAGCGGACTGGAAGCTGTCCGTACGATGAATGACTTGGGAATCAAGACCAAACACCCAATTGTC
GTTACCAATTGGACTAACGAGGAGGGGGCGCGCTTTGCACCTGCAAT**A**TTGGCCTCTGGCGTATTTGCAGGCGT**G**
CACACCTTAGAATATGCCTACGCCCGTAAGGACCCAGAAGGCAAATCTTTCGGTGATGAATTAAAGCGTATCGGC
TGGTTGGGCGATGAGGAAGTTGGTGCGCGTAAAATGCACGCATACTTTGAATACCACATCGAGCAAGGCCCAATT
CTTGAGGCGGAGAATAAGCAAATTGGCGTTGTCACTCATTGT**TTT**GGC**GGT**TGGTGGCTGGAATTTACTTTGACA
GGTCGCGAGGCTCATACCGGATCAACACCGATGGATATGCGTGTGAATGCGGGATTAGCGATGGCTCGTATTCTT
GAAATGGTCCAAACCGTTGCCATGGAGAACCAACCCGGTGCAGTGGGCGGCGTGGGGCAGATGTTTTTTAGCCCT
AACTCCCGCAACGTGTTACCGGGAAAAGTAGTTTTCACAGTAGACATCCGTTCCCCTGATCAAGCAAAGCTGGAT
GGAATGCGCGCACGCATTGAGGCAGAAGCGCCAAAAATTTGTGAGCGTCTGGGTGTCGGATGCTCGATTGAAGCA
GTA**TGT**CAC**GCG**GACCCAGTAACCTTTGACCCTAAACTTGTAGAGACAGTTCGTGGGGCGGCAGAGAAATTAGGG
TATTCGCATATGAACTTAGTTTCTGGCGCCGGACACGACGCCTGCTGGGCGGCTAAGGTTGCACCTACGACAATG
ATCATGTGTCCTTGTGTGGGCGGACTTTCGCATAATGAGGCGGAAGATATTTCTCGTGAATGGGCAGCAGCCGGT
GCAGACGTCCTTTTTCATGCTGTTTTGGAAACCGCTGAGATTGTGGAGTGA

**SmLcar_M141I**

ATGGCGGCGCCTGGTGAAAACCGTCGTGTAAATGCCGATCGCTTATGGGACTCGCTGATGGAAATGGCGAAAATT
GGCCCTGGAGTTGCGGGAGGTAACAACCGCCAGACCTTGACAGACGCTGACGGGGAAGGGCGCCGCCTGTTTCAA
TCCTGGTGTGAAGAAGCCGGCTTGTCGATGGGCGTCGATAAGATGGGCACCATGTTTCTTACACGCCCTGGAACT
GACCCCGACGCATTGCCAGTTCATATTGGCTCGCACCTGGATACTCAGCCTACCGGAGGAAAATTTGACGGCGTC
TTAGGTGTATTGAGCGGACTGGAAGCTGTCCGTACGATGAATGACTTGGGAATCAAGACCAAACACCCAATTGTC
GTTACCAATTGGACTAACGAGGAGGGGGCGCGCTTTGCACCTGCAAT**A**TTGGCCTCTGGCGTATTTGCAGGCGTA
CACACCTTAGAATATGCCTACGCCCGTAAGGACCCAGAAGGCAAATCTTTCGGTGATGAATTAAAGCGTATCGGC
TGGTTGGGCGATGAGGAAGTTGGTGCGCGTAAAATGCACGCATACTTTGAATACCACATCGAGCAAGGCCCAATT
CTTGAGGCGGAGAATAAGCAAATTGGCGTTGTCACTCATTGTCAAGGCCTGTGGTGGCTGGAATTTACTTTGACA
GGTCGCGAGGCTCATACCGGATCAACACCGATGGATATGCGTGTGAATGCGGGATTAGCGATGGCTCGTATTCTT
GAAATGGTCCAAACCGTTGCCATGGAGAACCAACCCGGTGCAGTGGGCGGCGTGGGGCAGATGTTTTTTAGCCCT
AACTCCCGCAACGTGTTACCGGGAAAAGTAGTTTTCACAGTAGACATCCGTTCCCCTGATCAAGCAAAGCTGGAT
GGAATGCGCGCACGCATTGAGGCAGAAGCGCCAAAAATTTGTGAGCGTCTGGGTGTCGGATGCTCGATTGAAGCA
GTAGGTCACTTCGACCCAGTAACCTTTGACCCTAAACTTGTAGAGACAGTTCGTGGGGCGGCAGAGAAATTAGGG
TATTCGCATATGAACTTAGTTTCTGGCGCCGGACACGACGCCTGCTGGGCGGCTAAGGTTGCACCTACGACAATG
ATCATGTGTCCTTGTGTGGGCGGACTTTCGCATAATGAGGCGGAAGATATTTCTCGTGAATGGGCAGCAGCCGGT
GCAGACGTCCTTTTTCATGCTGTTTTGGAAACCGCTGAGATTGTGGAGTGA
